# Single-cell RNA-sequencing of differentiating iPS cells reveals dynamic genetic effects on gene expression

**DOI:** 10.1101/630996

**Authors:** Anna SE Cuomo, Daniel D Seaton, Davis J McCarthy, Iker Martinez, Marc Jan Bonder, Jose Garcia-Bernardo, Shradha Amatya, Pedro Madrigal, Abigail Isaacson, Florian Buettner, Andrew Knights, Kedar Nath Natarajan, HipSci Consortium, Ludovic Vallier, John C Marioni, Mariya Chhatriwala, Oliver Stegle

## Abstract

Recent developments in stem cell biology have enabled the study of cell fate decisions in early human development that are impossible to study *in vivo*. However, understanding how development varies across individuals and, in particular, the influence of common genetic variants during this process has not been characterised. Here, we exploit human iPS cell lines from 125 donors, a pooled experimental design, and single-cell RNA-sequencing to study population variation of endoderm differentiation. We identify molecular markers that are predictive of differentiation efficiency, and utilise heterogeneity in the genetic background across individuals to map hundreds of expression quantitative trait loci that influence expression dynamically during differentiation and across cellular contexts.

## Introduction

The early stages of human embryogenesis involve dramatic and dynamic changes in cellular states. However, the extent to which an embryo’s genetic background influences this process has only been determined in a small number of special cases linked to rare large-effect variants that cause developmental disorders. This lack of information is critical - it can provide a deep understanding of how genetic heterogeneity is tolerated in normal development, when controlling the expression of key genes is vital. Additionally, with cellular reprogramming becoming an increasingly used tool in molecular medicine, understanding how inter-individual variability effects such differentiations is key.

Critically, recent technological developments have begun to facilitate such studies *in vitro*. In particular, the generation of population-scale collections of human induced pluripotent stem cells (iPSCs) [1,2] has allowed for assessing regulatory genetic variants in pluripotent [1,2] as well as in differentiated cells [3–5]. In addition, the rapid developments in single-cell RNA-seq now allow for assessing the molecular impact of genetic variability in a continuous manner across early human development.

Here, we use a pooled cell differentiation assay to study endoderm differentiation across a set of 125 human iPSC lines, profiling changes in gene expression via single-cell RNA-sequencing at 4 developmental timepoints [6]. Our study not only allows discovery of hundreds of novel expression Quantitative Trait Loci (eQTL) that vary across differentiation, but also enables the uncovering of genetic variants that impact the rate at which a cell line differentiates. Finally, we generalise approaches from studies of the interaction between genotype and environment (GxE) by leveraging the single-cell resolution of our study to investigate the interplay between genetic factors and cellular states.

### Population-scale single-cell profiling of differentiating iPS cells

We considered a panel of well-characterized human iPSC lines derived from 125 unrelated donors from the Human Induced Pluripotent Stem Cell initiative (HipSci) collection [1]. In order to increase throughput and mitigate the effects of batch variation, we exploited a novel pooled differentiation assay, combining sets of four to six lines in one well prior to differentiation (28 differentiation experiments performed in total; hereon “experiments”; Fig. 1A, **S1, S2**). Cells were collected at four differentiation time points (iPSC; one, two and three days post initiation - hereon day0, day1, day2 and day3) and their transcriptomes were assayed using full-length RNA-sequencing (Smart-Seq2 [7]) alongside the expression of selected cell surface markers using FACS (TRA-1-60, CXCR4; **Fig. S3, S4; Methods**). Following quality control (QC), 36,044 cells were retained for downstream analysis, across which 11,231 genes were expressed (**Fig. S5; Methods**). Exploiting that each cell line’s genotype acts as a unique barcode, we demultiplexed the pooled cell populations, enabling identification of the cell line of origin for each cell (similar to [8]; **Methods**). At each time point, cells from between 104 and 112 donors were captured, with each donor being represented by an average of 286 cells (after QC, **Fig. S2; Tables S1, S2; Methods**). The success of the differentiation protocol was validated using canonical cell-surface marker expression: consistent with previous studies [9], an average of 72% cells were TRA-1-60(+) in the undifferentiated state (day0) and an average of 49% of cells were CXCR4(+) three days post differentiation (day3; **Fig. S3**).

**Figure 1.**
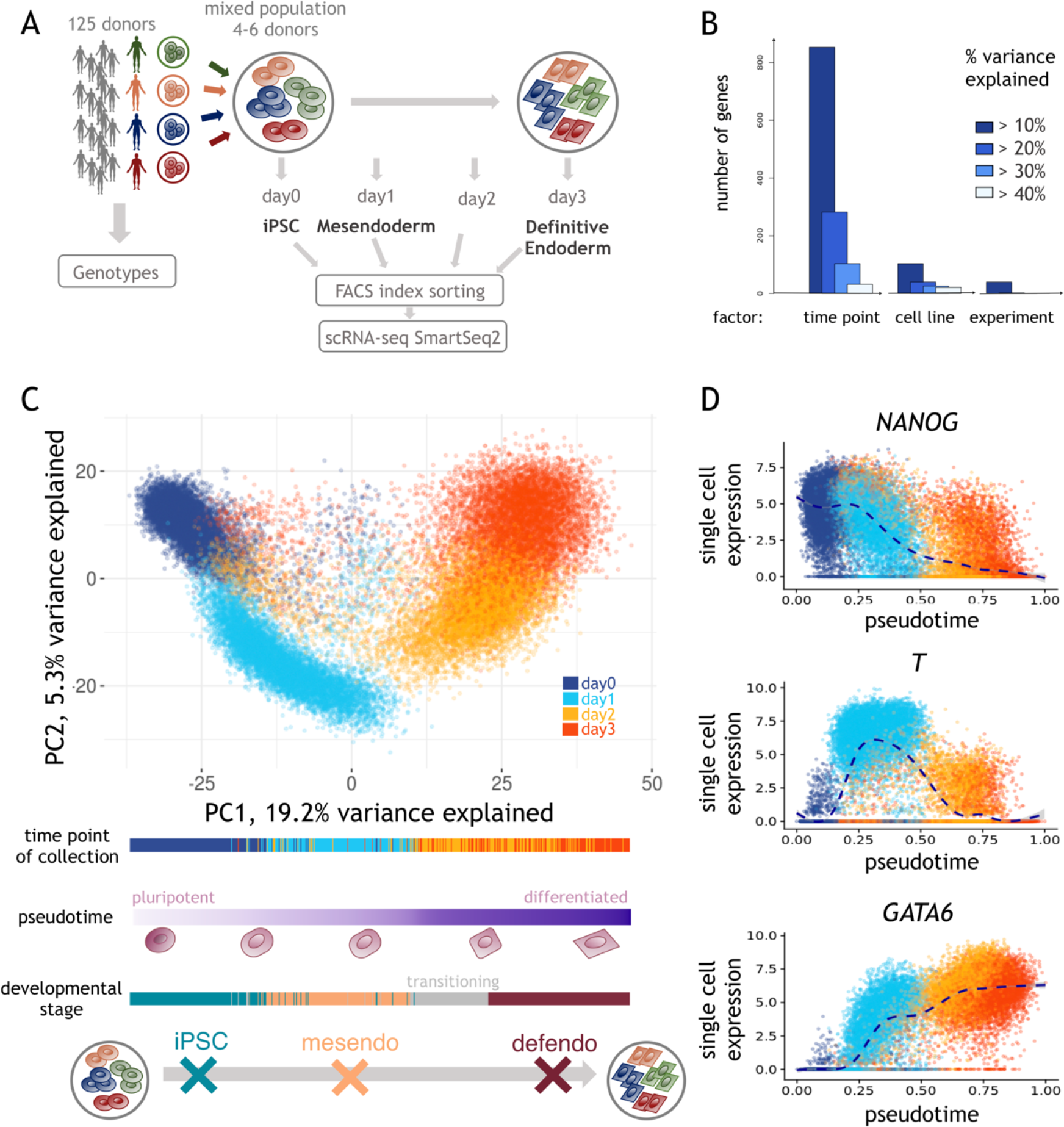
Single-cell endoderm differentiation of pooled iPSC lines. (**A**) Overview of the experimental design. iPSC lines from 125 genotyped donors were pooled in sets of 4-6, across 28 experiments, followed by differentiation towards definitive endoderm. Cells were sampled every 24 hours (**Methods**) and molecularly profiled using scRNA-seq and FACS. (**B**) Variance component analysis of 4,546 highly variable genes, using a linear mixed model fit to individual genes to decompose expression variation into time point of collection, cell line and experimental batch (**Methods**). (**C**) **Top:**Principal component analysis of gene expression profiles for 36,044 QC-passing cells. Cells are coloured by the time point of collection. **Bottom:** Cells are ordered by pseudotime, defined as the first principal component (PC1). From left to right, cells transition from a pluripotent state to definitive endoderm. (**D)**Single cell expression (y axis) of selected markers for each developmental stage, spanning iPSC (*NANOG*), mesendo (*T*), and defendo (*GATA6*) stages, plotted along pseudotime (x axis).

Variance component analysis across all genes (using a linear mixed model; **Methods**) revealed the time point of collection as the main source of variation, followed by the cell line of origin and the experimental batch (Fig. 1B). Consistent with this, the first Principal Component (PC) was strongly associated with differentiation time (Fig. 1C, **S6; Methods**), motivating its use to order cells by their differentiation status (hereafter “pseudotime”, Fig. 1C). Alternative pseudotime inference methods yielded similar orderings (**Fig. S7; Methods**).

Critically, the expected temporal expression dynamics of marker genes that characterise endoderm differentiation was captured by the ordering of cells along the inferred pseudotime (Fig. 1D). Exploiting these markers of differentiation progress and pseudotime, we assigned 28,971 cells (~80%) to one of three canonical stages of endoderm differentiation: iPSC, mesendoderm (mesendo) and definitive endoderm (defendo) (Fig. 1C, **S8; Methods**). A smaller fraction of cells (N = 7,073) could not be confidently assigned to a canonical stage of differentiation; these cells were heavily enriched for those collected at day2, when rapid changes in molecular profiles are expected, reflecting a transitional population of cells.

### Pseudo-temporal ordering yields stage-specific eQTL

Motivated by the observation that a substantial fraction of variability in gene expression was explained by cell-line effects (Fig. 1B), we tested for associations between common genetic variants and gene expression at the three defined stages of cell differentiation (Fig. 1C). Briefly, for each donor, experimental batch, and differentiation stage, we quantified each gene’s average expression level (**Methods**), before using a linear mixed model to test for *cis* eQTL, adapting approaches used for bulk RNA-seq profiles (+/− 250kb, MAF > 5% [1]; **Methods**). In the iPSC population (day0), this identified 1,833 genes with at least one eQTL (denoted eGenes; FDR < 10%; 10,840 genes tested; **Table S3**). To validate our approach, we also performed eQTL mapping using deep bulk RNA-sequencing data from the same set of iPSC lines (“iPSC bulk”; 10,736 genes tested), yielding consistent eQTL (~70% replication of lead eQTL effects; nominal P < 0.05; **Methods; Table S4**).

Analogously, we mapped eQTL in the mesendo and defendo populations, yielding 1,702 and 1,342 eGenes respectively. For comparison, we also performed eQTL mapping in cells collected on day1 and day3 – the experimental time points commonly used to identify cells at mesendo and defendo stages [6]. Interestingly, this approach identified markedly fewer eGenes (1,181 eGenes at day1, and 631 eGenes at day3), demonstrating the power of using the single-cell RNA-seq profiles to define relatively homogeneous differentiation stages in a data-driven manner (Fig. 2B, **S9; Methods; Table S5**).

**Figure 2.**
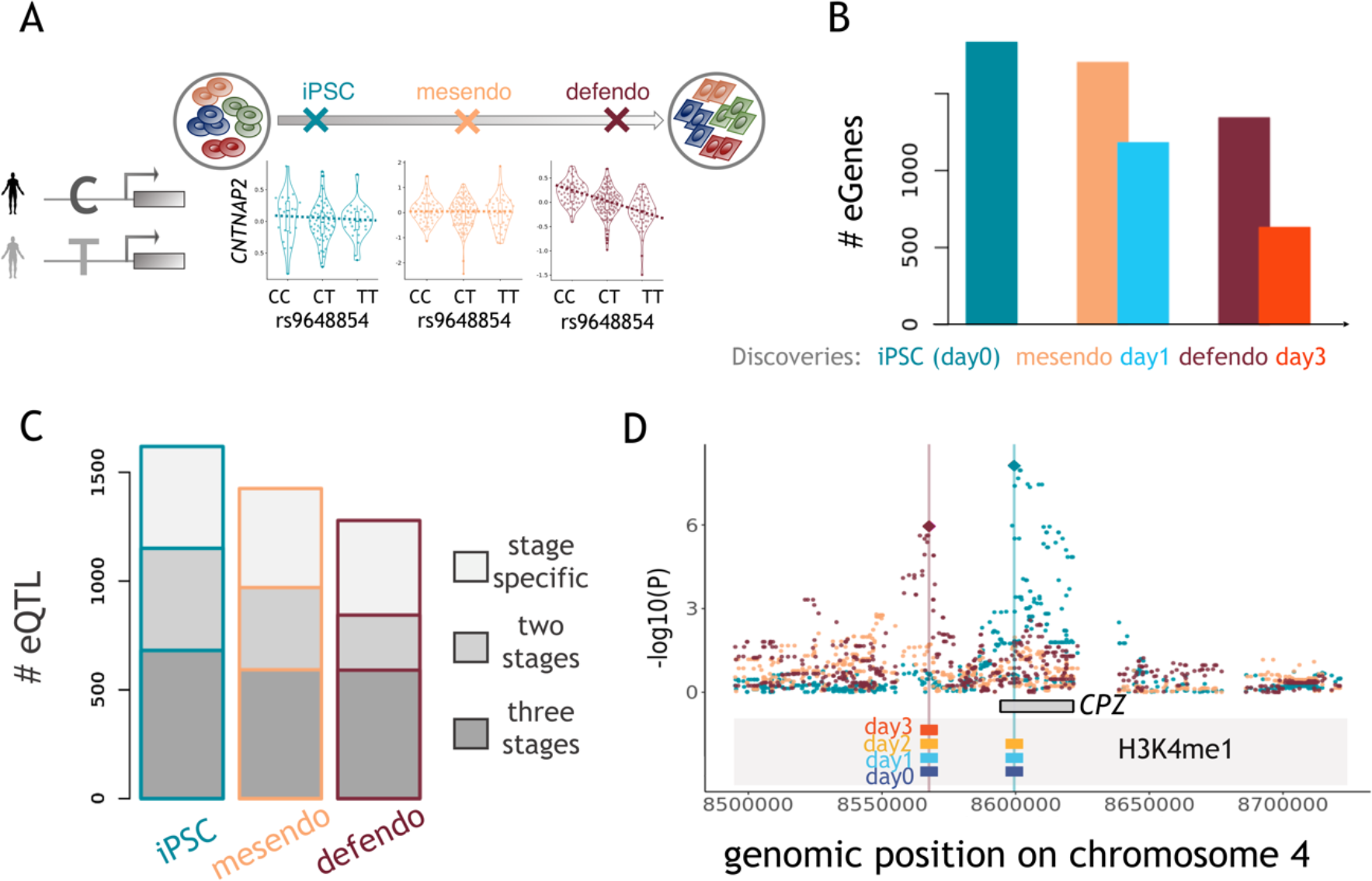
Mapping single-cell eQTL in each developmental stage. (**A**) Illustration of the single cell eQTL mapping strategy at different stages of differentiation. Shown is an example of an eQTL that is specific to the defendo stage. Boxplots of gene expression stratified by the allelic state of rs9648854 at each stage, showing an association between rs9648854 and *CNTNAP2* expression at the defendo stage, but not at earlier stages. (**B**) Comparison of eQTL mapping using different strata of all cells. Stage definition based on pseudotime ordering increases the number of detectable eQTL, compared to using the time point of collection. Bars represent number of eGenes (genes with at least one eQTL, at FDR < 10%). (**C**) Proportion of eQTL that are specific to a single stage, shared across two stages, or observed across all stages (sharing defined as a lead eQTL variant at one stage with nominal significant effects P < 0.05 and consistent direction at another stage). (**D**) A lead switching event consistent with epigenetic remodelling. The overlap of H3K4me1 with the eQTL SNPs across differentiation time points is indicated by the coloured bars.

Profiling multiple stages of endoderm differentiation allowed us to assess at which stage along this process individual eQTL can be detected. We observed substantial regulatory and transcriptional remodelling upon iPS differentiation to definitive endoderm, with over 30% of eQTL being specific to a single stage (Fig. 2A, 2C; **Methods**). Our differentiation time course covers developmental stages that have never before been accessible to genetic analyses of molecular traits. Consistent with this, 349 of our eQTL variants at the mesendo and defendo stages have not been reported in either a recent iPSC eQTL study based on bulk RNA-seq [10], or in a compendium of eQTL identified from 49 tissues as part of the GTEx project [11] (linkage disequilibrium, LD: r^2^ < 0.2; **Methods; Table S3**).

In addition to these novel eQTL, we identified lead switching events for 155 eGenes, that is distinct lead eQTL variants for the same gene at different stages of differentiation (LD: r^2^ < 0.2; **Methods**). To investigate the potential regulatory role of such variants, we examined whether the corresponding genetic loci also featured changes in histone modifications during differentiation. Specifically, we used ChIP-Sequencing to profile five histone modifications associated with gene and enhancer usage (H3K27ac, H3K4me1, H3K4me3, H3K27me3, H3K36me3) in hESCs that were differentiated (using the same protocol employed above) towards endoderm and measured at equivalent time points (i.e. day0, day1, day2, day3; **Methods**). Intriguingly, for 20 of the lead switching events, we observed corresponding changes in the epigenetic landscape (stage-specific lead variants overlap with stage-specific changes in histone modification status), suggesting a direct mode of action (Fig. 2D).

### eQTL variants and early molecular markers are predictive of differentiation efficiency

Previous studies have demonstrated that iPSC lines vary in their capacity to differentiate [12]. As a measure of differentiation efficiency in our experiments, we used average pseudotime on day3, and observed significant variation across cell lines, which was consistent across replicate differentiations of the same cell line (Fig. 3A). Exploiting the scale of our study and the pooled experimental design, we set out to identify genetic and molecular markers of differentiation efficiency that are accessible prior to differentiation **(Methods**).

**Figure 3.**
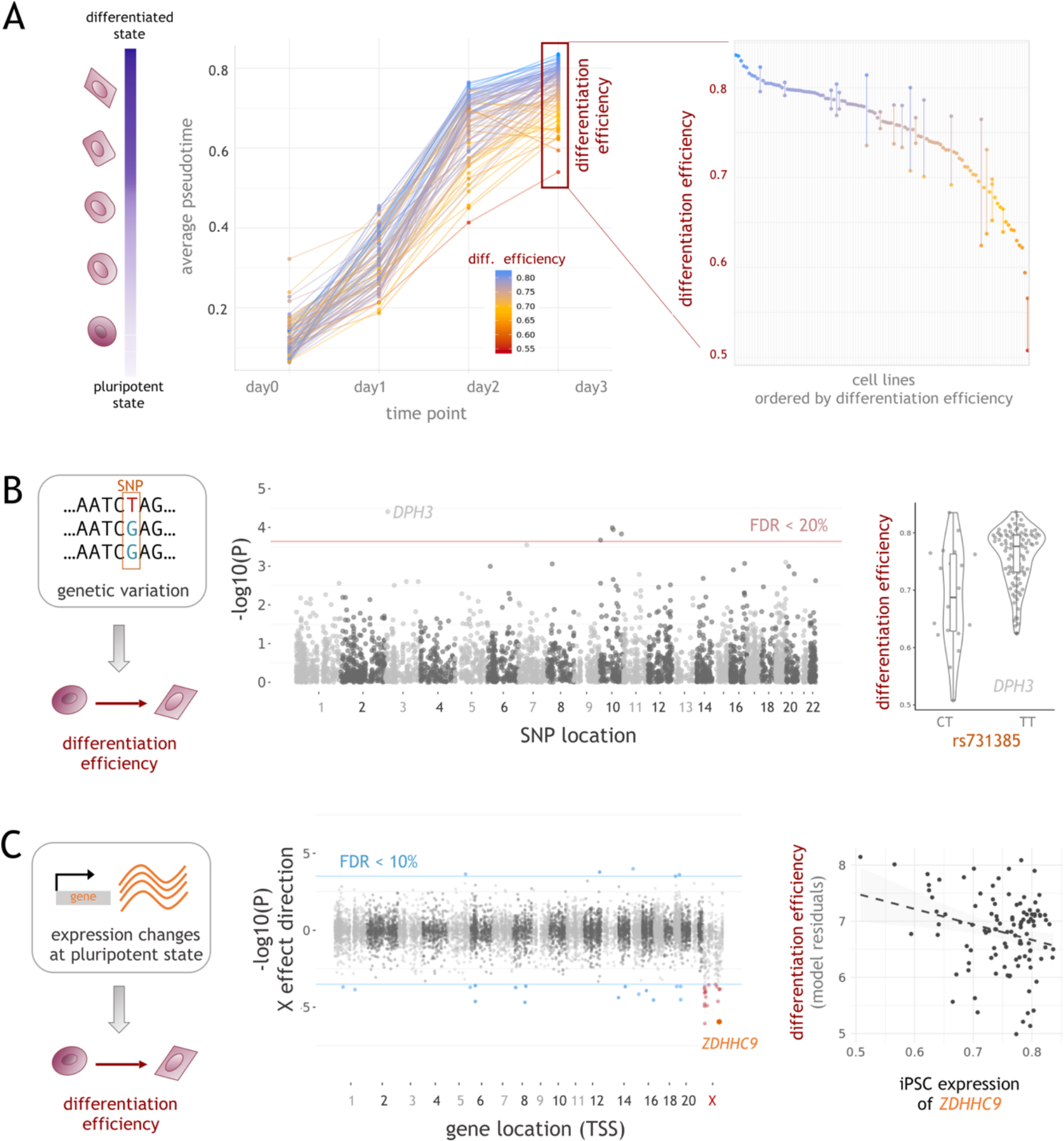
Identification of molecular markers for differentiation efficiency. (**A**) Variation in differentiation efficiency across cell lines. **Left:** Differentiation progress over time, showing trajectories for 98 cell lines, coloured by differentiation efficiency. Shown are 98 cell lines with sufficient data at all time points (out of 126, more than 10 cells). Differentiation efficiency of a cell line was defined as the average pseudotime across all cells on day 3. **Right**: Differentiation efficiency across cell lines (points), and consistency of individual cell lines differentiated in multiple experiments (vertical bars). (**B**) Effects of genetic variation on differentiation efficiency. **Left:** schematic. **Center:** Manhattan plot displaying negative log P values for association tests between 4,422 lead eQTL variants and differentiation efficiency. Highlighted is an association for an eQTL variant for *DPH3*. Horizontal red line denotes FDR = 20% (Benjamini-Hochberg adjusted). **Right:** Boxplot displaying differentiation efficiency for 125 individuals stratified by the allelic state of rs73138519 (mesendo eQTL for *DPH3)*, which is associated with decreased differentiation efficiency (**Methods**). (**C**) Associations between iPSC gene expression levels and differentiation efficiency. **Left:** schematic. **Center:** Genome-wide analysis to identify markers of differentiation efficiency, considering iPSC gene expression levels. Displayed are negative log P values signed by the direction of the effect. Horizontal blue lines denote FDR = 10% (Benjamini-Hochberg adjusted). Autosomal genes with significant associations are shown in blue; X chromosome genes with significant associations are shown in red. **Right:** Scatter plot between gene expression in the iPS state and differentiation efficiency for the X chromosome gene *ZDHHC9*.

First, we considered the set of 4,422 eQTL lead variants at any of the three developmental stages and tested each variant for association with differentiation efficiency (Fig. 3B; using a linear mixed model; **Methods**). This identified 5 eQTL variants at a lenient false discovery rate threshold (FDR 20%; Fig. 3B, **Table S6**). The most significant associations were observed with eQTL variants *for DPH3* (P = 3.9e-5) and *H2AFY2* (P = 1e-4). Loss of *DPH3* results in an embryonic lethal phenotype in mice [13], while the effect direction of the eQTL variant for *H2AFY2* was consistent with observations that knockdown of this gene inhibits endoderm differentiation of human iPSCs *in vitro* [14]. In order to further investigate these associations, we used staining for the percentage of CXCR4+ as an independent measure of differentiation efficiency [15]. CXCR4+ staining data on the same lines enabled replication of 3/5 of these associations (P < 0.05; one-tailed test). We also performed an additional set of differentiations in iPSC lines derived from individuals that were not part of the variant discovery, selected based on genotype at the *DPH3* eQTL locus (n = 20). While the direction of effect was consistent, the association was not statistically significant (P = 0.24), likely reflecting low power at this sample size. Collectively, these results indicate that our approach can reveal genetic determinants of *in vitro* differentiation efficiency.

Having identified genetic markers associated with differentiation capacity we next asked whether the average expression level of genes at the iPSC stage could represent molecular markers of differentiation efficiency. This revealed 38 associations (FDR 10%, 11,231 genes tested; **Table S7**), 15 of which were also observed when using independent bulk RNA-seq data from the same cell lines (replication defined as nominal P < 0.05; **Table S7; Methods**). As an example, the expression of *ZDHHC9* in iPSCs was negatively associated with differentiation efficiency (Fig. 3C). Furthermore, *ZDHHC9* is one of 17 differentiation-associated genes located on the X chromosome, reflecting a significant enrichment of X chromosome genes (24.5-fold enrichment, P = 8×10^−16^, Fisher’s exact test). Higher expression of these genes was associated with reduced differentiation efficiency (Fig. 3C; **Methods**). The majority of these associations persisted when limiting the analysis to female lines (14/17 at P < 0.05), indicating variation beyond differences between sexes. These results are consistent with previous observations that X chromosome reactivation is a marker of poor differentiation capacity of iPSCs in general [16,17]. Finally, we note that the set of associated genes located on other chromosomes included genes with known roles in iPSC differentiation, such as *TBX6* [18].

### Discovery of dynamic eQTL across iPSC differentiation

The availability of large numbers of cells per donor across the differentiation trajectory enabled the analysis of dynamic changes of eQTL strength at fine-grained resolution. Using a sliding-window approach (25% cells in each window, sliding along pseudotime by a step of 2.5% cells), we assessed how the joint set of 4,422 eQTL lead variants (4,470 SNP-gene pairs) discovered at the iPSC, mesendo, and defendo stages were modulated by developmental time. To do this, we reassessed each eQTL in each window, recording a SNP effect size per window (**Methods)**. As a complementary approach, we also took advantage of the full length transcript sequencing to measure allele-specific expression (ASE) in each window (Fig. 4A **top panel; Methods**). Here, in each window, we quantified the deviation from 0.5 of the expression of the minor allele at the eQTL (ratio of reads phased to eQTL variants, **Methods**). Both methods result in a measure of the varying strength of genetic effects along development, or genetic effect dynamics. Reassuringly, the two approaches were highly consistent across pseudotime (Fig. 4A, **S10**).

**Figure 4.**
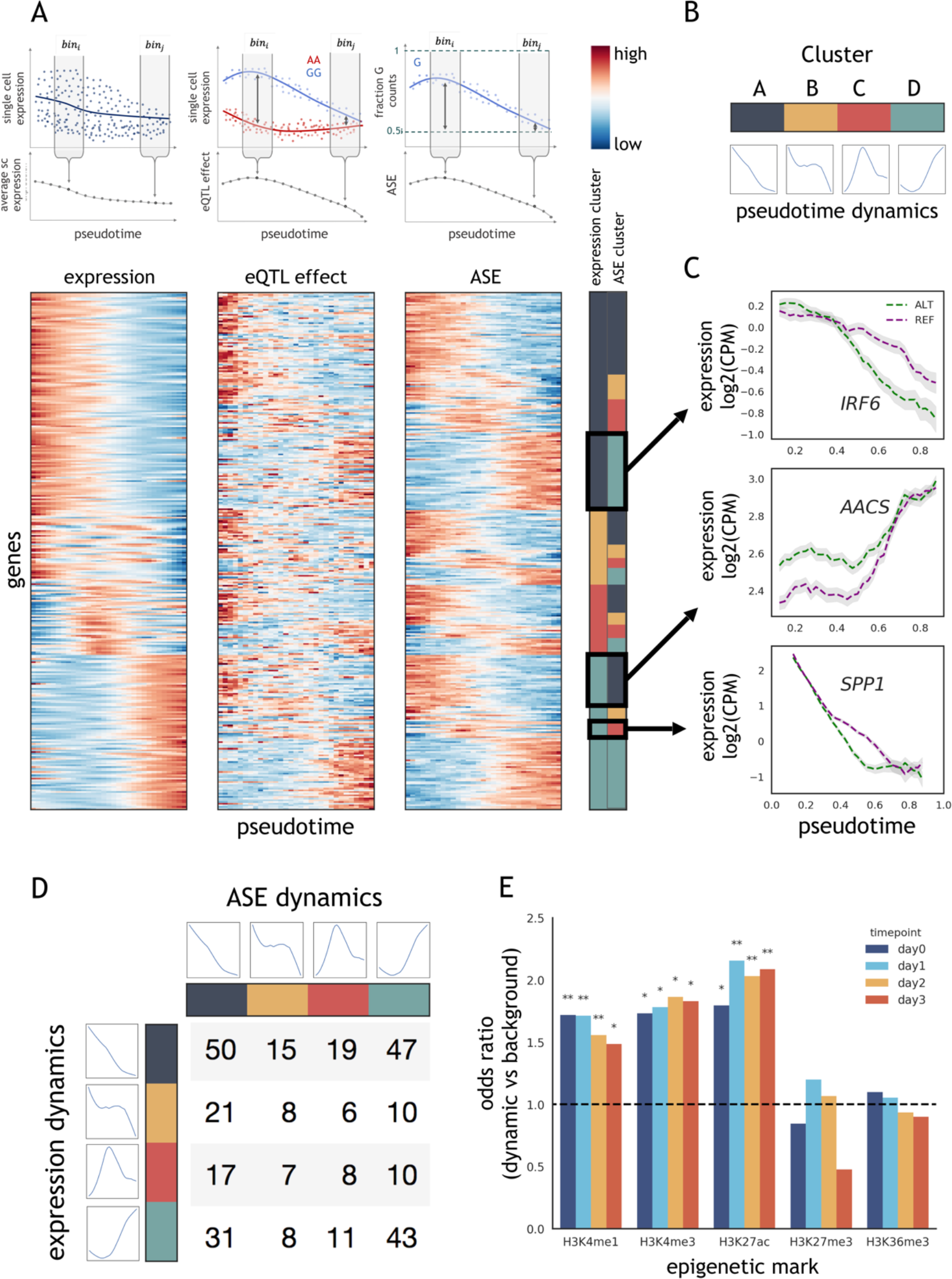
eQTL dynamics during differentiation. (**A**) Combined analysis of the gene expression, ASE, and eQTL dynamics across pseudotime. **Upper panels:** Schematic of sliding window approach. Cells are binned according to pseudotime groups, to quantify average expression, perform an eQTL analysis, and quantify average ASE (each bin includes 25% of cells, binned at increments of 2.5%). **Lower panels:** clustered heatmap of expression levels, eQTL effects, and ASE across pseudotime for the top 311 genes with the strongest dynamic QTL effects (FDR < 1%; out of 785 at FDR < 10%; **Methods**). For each gene, the expression and the ASE dynamics were jointly grouped using clustering analysis, with 4 clusters. The membership of gene expression and ASE dynamics of these 4 clusters is indicated by colours in the right-hand panel. Values in all heatmaps are z-score normalised by row. For ASE, average ASE values are plotted such that red indicates highest deviation from 0.5. (**B**) Summary of the identified cluster dynamics, displaying the average dynamic profile of each cluster, computed as the average across z-score normalized gene expression/ASE profiles. (**C**) Exemplars of the dynamic gene expression and dynamic genetic effects clusters shown in **A**. Shaded regions indicate standard error (+/− 1 SEM; **Methods**). (**D**) Number of genes categorized by the combination of expression and ASE cluster from **A**. Average dynamics of expression clusters (rows) and ASE clusters (columns) as in **B** are shown. (**E**) Overlap of dynamic eQTL variants from **A** with histone marks. The odds ratio compared to the background of all other eQTL variants is shown (*P < 0.01; **P < 1×10^−4^; Fisher’s exact test).

To formally test for eQTL effects that change dynamically across differentiation (dynamic QTL), we tested for associations between pseudotime and the genetic effect size (defined based on ASE; likelihood ratio test, considering linear and quadratic pseudotime), uncovering a total of 785 time dynamic eQTL (FDR < 10%; **Methods**), including a substantial fraction of eQTL that were not stage-specific (**Table S3**). This complements our earlier analysis, which identified substantial stage-specific effects (Fig. 2A, 2C), by identifying subtle changes in the relationship between genotype and phenotype during differentiation. To further explore this set of genes, we clustered eQTL jointly based on the relative gene expression dynamics (global expression changes along pseudotime, quantified in sliding windows as above, **Methods**), and on the genetic effect dynamics (Fig. 4A; **Methods**). This identified four basic dynamic patterns (Fig. 4B): sharply decreasing (cluster A), gradually decreasing (cluster B), transiently increasing (cluster C), and gradually increasing (cluster D). As expected, stage-specific eQTL were grouped together in particular clusters (e.g. defendo specific eQTL in cluster D; **Fig. S11**). Notably, the gene expression dynamics and the eQTL dynamics tended to be distinct, demonstrating that gene expression level is not the primary mechanism governing variation in genetic effects. In particular, genetic effects were not most pronounced when gene expression was high (Fig. 4C, 4D).

Distinct combinations of expression and eQTL dynamics result in different patterns of allelic expression. This is illustrated by the mesendoderm-specific eQTL for *SPP1*. Overall expression of *SPP1* decreases during differentiation, but expression of the alternative allele is repressed more quickly than that of the reference allele (Fig. 4C). This illustrates how *cis* regulatory sequence variation can modulates the timing of expression changes in response to differentiation, similar to observations previously made in *C. elegans* using recombinant inbred lines [19]. In other cases, the genetic effect coincides with high or low expression, for example in the cases of *IRF6* and *AACS* (Fig. 4C). These examples illustrate how genetic variation is intimately linked to the dynamics of gene regulation.

We next asked whether dynamic eQTL were located in specific regulatory regions. To do this, we evaluated the overlap of the epigenetic marks defined using the hESC differentiation time series with the dynamic eQTL (Fig. 4D, **S12**). This revealed an enrichment of dynamic eQTL in H3K27ac, H3K4me1 (i.e. enhancer marks), and H3K4me3 (i.e. promoter) marks compared to non-dynamic eQTL (i.e. eQTL that we identified but did not display dynamic changes along pseudotime, Fig. 4D), consistent with these SNPs being located in active regulatory elements.

### Cellular environment modulates genetic effects on expression

Whilst differentiation was the main source of variation in the dataset, single cell RNA-seq profiles can be used to characterize cell-toll-cell variation across a much wider range of cell state dimensions [20–22]. We identified sets of genes that varied in a co-regulated manner using clustering (affinity propagation; 8,000 most highly expressed genes; **Table S8**; **Methods**), which identified 60 modules of co-expressed genes. The resulting modules were enriched for key biological processes such as cell differentiation, cell cycle state (G1/S and G2/M transitions), respiratory metabolism, and sterol biosynthesis (as defined by Gene Ontology annotations; **Table S9**). These functional annotations were further supported by transcription factor binding (e.g. enrichment of SMAD3 and E2F7 targets in the differentiation and cell cycle modules, respectively (**Table S10, S11**)). Additionally, expression of the cell differentiation module (cluster 6; **Table S9**) was correlated with pseudotime, as expected (R = 0.62; **Fig. S7**).

Using the same ASE-based interaction test as applied to identify dynamic QTL, reflecting ASE variation across pseudotime (Fig. 4; **Methods**), we assessed how the genetic regulation of gene expression responded to these cellular contexts. Briefly, we tested for genotype by environment (GxE) interactions using a subset of four co-expression modules as markers of cellular state, while accounting for pseudotime (Fig. 5A; **Methods**). These four co-expression modules were annotated based on GO term enrichment, and taken as markers representing cell cycle state (G1/S and G2/M transitions) and metabolic pathway activity (respiratory metabolism and sterol biosynthesis; **Methods**). This approach extends previous work using ASE to discover GxE interactions [23,24], taking advantage of the resolution provided by single-cell data. We identified 686 eQTL that had an interaction effect with at least one factor (Fig. 5B; FDR < 10%), with many of these effects being orthogonal to the effects of differentiation. Indeed, 394 genes had no association with pseudotime, but responded to at least one other factor. Conversely, of the 785 dynamic eQTL, 292 were also associated with other factors, while 493 were associated only with pseudotime (Fig. 5B, **S13; Tables S13; Methods**).

**Figure 5.**
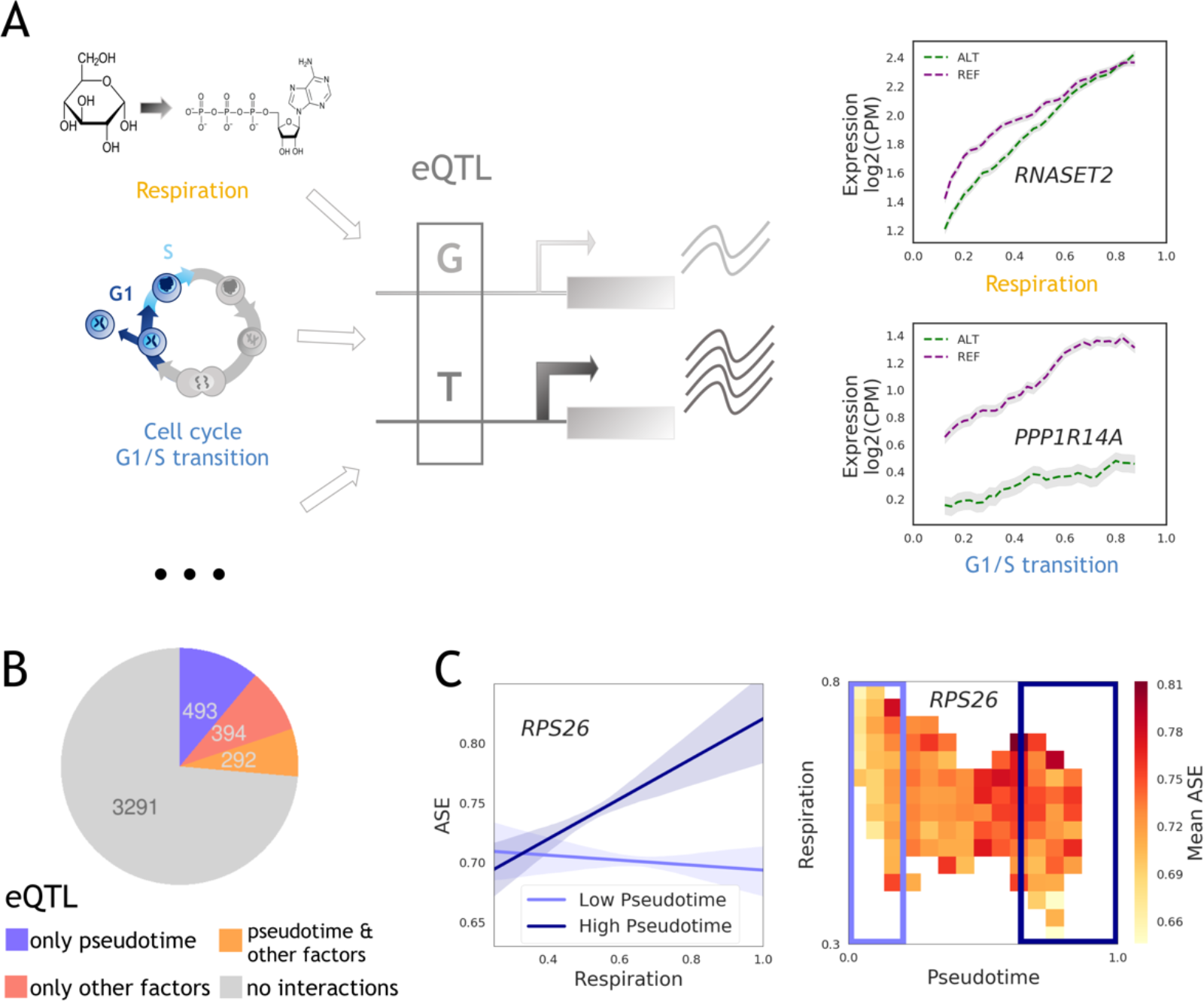
Allele-specific expression reveals interactions with fundamental cellular processes. (**A**) Illustration of eQTL affected by cellular context. **Left:** Schematic of cellular contexts affecting a regulatory element containing an eQTL SNP, and thus affecting allele-specific expression. **Right:** Allele-specific expression variation for two exemplar eQTL SNPs that tag cancer GWAS variants and display GxE interactions (FDR < 10%). The eQTL for *RNASET2* (rs2247315) tags a risk variant for basal cell carcinoma, and is responsive to cellular respiration, while that for *PPP1R14A* (rs12608912) tags a risk variant for prostate cancer and is responsive to the cell cycle G1/S transition (**Table S12**). Cellular contexts for each cell were inferred by GO annotations of coexpression modules (**Methods**). Shaded regions indicate standard error (+/− 1 SEM; **Methods**). (**B**) Results summary: numbers of eQTL (from Fig.2; **Methods**) identified as displaying GxE interactions with pseudotime (purple), displaying GxE interactions with other cellular contexts but not with pseudotime, (after appropriately accounting for pseudotime, red), displaying GxE interactions with both pseudotime and at least one other cellular context (yellow), and displaying no GxE interactions at all (grey). Significance is assessed at FDR < 10%. (**C**) Higher order interaction example: an eQTL variant for *RPS26* (rs10876864) is affected by a GxExE higher order interaction with both pseudotime and respiration. This variant is also a risk variant for allergic disease and vitiligo. **Left panel:** Effects of respiration state on ASE for cells with low and high pseudotime. Lines shown are linear regressions with 95% confidence intervals for the 30% of cells with lowest and highest values for pseudotime. **Right panel:** Heatmap of averaged ASE for cells falling within the specified windows of pseudotime and respiration state. Only values for windows containing n > 30 cells are shown (n = 17,373 cells in total).

These interactions encompass regulatory effects on genes and SNPs with important functional roles. Specifically, 145 interaction eQTL variants overlap with variants previously identified in genome-wide association studies (GWAS, LD r^2^ > 0.8; **Methods; Table S12**), including seven risk variants for cancer (EFO term: EFO_0000311). For example, an eQTL for *RNASET2* shows sensitivity to cellular respiratory metabolic state (Fig. 5A). This eQTL SNP is in strong LD (r^2^ = 1.0) with a GWAS risk variant for basal cell carcinoma [25]. Furthermore, an eQTL for *PPP1R14A* showed sensitivity to the G1/S state, and is in LD (r^2^ = 0.81) with a GWAS risk variant for prostate cancer [26] (Fig. 5A). The onset of cancer affects cellular respiratory metabolism and cell cycle progression [27], raising the possibility that the effects of these variants are enhanced during oncogenesis. These examples illustrate the versatility of our single cell dataset and how it can provide regulatory information about variants in contexts beyond early human development.

Finally, we explored whether we could detect higher order interaction effects, where the genetic effect varies with a cellular state in different ways along differentiation, effectively testing for GxExE interactions. To this end, we fitted a linear model with fixed effects for differentiation and each of the factors, plus a combined term (factor x pseudotime, Fig. 5B, 5C; **Methods**). This identified 220 genes with significant higher order interactions between a genetic variant, differentiation, and at least one other factor (Fig. 5B, 5C, **S13; Table S13d**). One example is the eQTL for *RPS26*, whose ASE was sensitive to cellular respiration, but only late in differentiation (Fig. 5C). This eQTL variant (rs10876864) is a risk variant for allergic disease and vitiligo [28,29]. These results highlight the context-specificity of eQTL, and the power of scRNA-seq in dissecting this specificity within one set of experiments.

## Discussion

Our map of early endoderm differentiation across 125 individuals offers a unique and powerful tool for interrogating the role of genetic heterogeneity in early human development. We exploited this resource to identify hundreds of novel eQTL that act at tightly-defined time points during early differentiation, and at specific states, thus fully utilising the power of single-cell transcriptomics. Moreover, we used our map and an independent experimental validation assay to demonstrate that specific germline variants have the potential to alter the rate of differentiation.

More generally, this latter analysis elucidates the broad utility of our data for studying the role of genetic variation in regenerative medicine and normal development. In the case of definitive endoderm differentiation, the *in vitro* protocol is short and efficient, the molecular basis is relatively well understood, and the process is highly canalised [30]. However, most differentiation protocols are less well understood, less efficient, more variable, and require more time. Thus, we expect application of this approach in other contexts to expand our molecular understanding, improve protocol efficiency, and characterise the genetic component of differentiation across the spectrum of human development and cellular contexts.

## Methods

### Overview: pooled scRNA-seq profiling during endoderm differentiation

A total of 126 pluripotent stem cell (iPSC) lines derived from 125 donors as part of the HipSci project were considered for analysis (**Table S1**). Batches of 4-6 cell lines were co-cultured and grown as a mixed population for a total of 28 experiments, in 12 well plates. Cells were harvested immediately prior to the initiation of differentiation (day0; iPSCs), and at time points 1, 2, and 3 days post differentiation initiation (day1, day2, day3). Subsequently, single cells were sorted into 384 well plates. Cells were processed using Smart-seq2 for scRNA-seq with parallel FACS analysis of the markers TRA-1-60 and CXCR4 being performed for each cell. A subset of cell lines were assayed in more than one experiment (33 donors; **Table S1, S2; Fig. S2**). In addition to the differentiation of pools of cell lines by co-culture for scRNA-seq, cell lines were also differentiated individually and assayed by FACS for the percentage of CXCR4+ cells on day3, following the same protocol. These individual differentiations were performed in two phases. First, individual differentiations of cell lines included in the scRNA-seq experiments were performed in parallel with the single-cell experiments. Second, an independent set of differentiations of new cell lines (i.e. cell lines derived from individuals not represented in the first set of cell lines), selected by genotype in order to validate the genetic association with differentiation, were performed as separate experiments.

### Cell culture for maintenance and differentiation

Human iPSC lines were thawed for differentiation and maintained in Essential 8 (E8) media (LifeTech) according to the manufacturer's instructions. Prior to plating for differentiation, cells were passaged at least twice after thawing and always 3 - 4 days before plating for differentiation to ensure all the cell lines in each experiment were growing at a similar rate prior to differentiation. To plate for endoderm differentiation, cells were washed 1x with DPBS and dissociated using StemPro Accutase (Life Technologies, A1110501) at 37°C for 3 - 5 min. Colonies were fully dissociated through gentle pipetting. Cells were resuspended in MEF medium [6], passed through a 40μm cell strainer, and pelleted gently by centrifuging at 300xg for 5 min. Cells were re-suspended in E8 media and plated at a density of 15,000 cells per cm^2^ in gelatin/MEF coated plates [6,31] in the presence of 10 μM Rock inhibitor – Y27632 [10 mM] (Sigma, Cat # Y0503 - 5 mg). Media was replaced with fresh E8 free of Rock inhibitor every 24 hours post plating. Differentiation into definitive endoderm commenced 72 hours post plating as previously described [6]. The overall efficiency of the differentiation protocol was validated using reference lines with good and poor differentiation capacity, respectively (**Fig. S14**).

### Single cell preparation and sorting for scRNAseq

Cells were dissociated into single cells using Accutase and washed 1x with MEF medium as described above. For all subsequent steps, cells were kept on ice to avoid degradation. Approximately 1 x 10^6^ cells were re-suspended in PBS + 2% BSA + 2 mM EDTA (FACS buffer); BSA and PBS were nuclease-free. For staining of cell surface markers TRA-1-60 (BD560380) and CXCR4 (eBioscience 12-9999-42), cells were re-suspended in 100 μL of FACS buffer with enough antibodies to stain 1 x 10^6^ cells according to the manufacturer’s instructions, and were placed on ice for 30 min. Cells were protected from light during staining and all subsequent steps. Cells were washed with 5 mL of FACS buffer, passed through a 35 μM filter to remove clumps, and re-suspended in 300 μL of FACS buffer for live cell sorting on the BD Influx Cell Sorter (BD Biosciences). Live/dead marker 7AAD (eBioscience 00-6993) was added immediately prior to analysis according to the manufacturer's instructions and only living cells were considered when determining differentiation capacities. Living cells stained with 7AAD but not TRA-1-60 or CXCR4 were used as gating controls. Data for TRA-1-60 and CXCR4 staining were available for 31,724 cells, of the total 36,044. Single-cell transcriptomes of sorted cells were assayed as follows: reverse transcription and cDNA amplification was performed according to the SmartSeq2 protocol [7], and library preparation was performed using an Illumina Nextera kit. Samples were sequenced using paired-end 75bp reads on an Illumina HiSeq 2500 machine (one lane of sequencing per 384 well plate).

### Genotyping

iPS cell lines were genotyped as previously described [1], using the Illumina HumanCoreExome-12 Beadchip. Genotypes were called using GenomeStudio (Illumina, CA, USA), following independent imputation using IMPUTE2 v2.3.1 [32] and phasing using SHAPEIT v2.r790 [33]. Imputation was performed based on a joint reference panel of haplotypes from the UK10K cohorts and 1000 Genomes Phase 1 data [33,34]. Single-sample VCFs were merged and subsequent QC was performed using Genotype Harmonizer [35] and BCFtools. Variants with INFO score lower than 0.4 were excluded from further analysis.

### Demultiplexing donors from pooled experiments

Assignment of cells to donors was performed using Cardelino [36]. Briefly, Cardelino estimates the posterior probability of a cell originating from a given donor based on common variants in scRNA-seq reads, while employing a beta binomial-based Bayesian approach to account for technical factors (e.g. differences in read depth, allelic drop-out, and sequencing accuracy). For this assignment step, we considered a larger set of n = 490 HipSci lines with genotype information, which included the 126 lines used in this study. A cell was assigned to a donor if the model identified the match with posterior probability > 0.9, requiring a minimum of 10 informative variants for assignment. Cells for which the donor identification was not successful were not considered further. Across the full dataset 99% of cells that passed RNA QC steps (below) were successfully assigned to a donor.

### scRNA-seq quality control and processing

Adapters of raw scRNA-seq reads were trimmed using Trim Galore! [37–39], using default settings. Trimmed reads were mapped to the human reference genome build 37 using STAR [40] (version: 020201) in two-pass alignment mode, using the default settings proposed by the ENCODE consortium (STAR manual). Gene-level expression quantification was performed using Salmon [41] (version: 0.8.2), using the “--seqBias”, “–gcBias” and “VBOpt” options using ENSEMBL transcripts (built 75) [42]. Transcript-level expression values were summarized at a gene level (estimated counts per million (CPM)) and quality control of scRNA-seq data was performed with the *scater* Bioconductor package in R [43]. Cells were retained for downstream analyses if they had at least 50,000 counts from endogenous genes, at least 5,000 genes with non-zero expression, less than 90% of counts came from the 100 highest-expressed genes, less than 15% of reads mapping to mitochondrial (MT) genes, they had a Salmon mapping rate of at least 60%, and if the cell was successfully assigned to a donor (**Fig. S15**). Dead cells as identified based on 7AAD staining were discarded. Size factor normalization of counts was performed using the *scran* Bioconductor package in R [44]. Expressed genes with an HGNC symbol were retained for analysis, where expressed genes in each batch of samples were defined based on i) raw count > 100 in at least one cell prior to QC and ii) average log2(CPM+1) > 1 after QC. Normalized CPM data were log transformed (log2(CPM+1)) for all downstream analyses. The joint dataset was investigated for outlying cell lines or experimental batches, which identified no clear groups of outlying cells (**Fig. S16, S17**).

As a final QC assessment, we considered possible differences between cell lines from healthy and diseased donors. In particular, a subset of 11 cell lines were derived from neonatal diabetes patients, and differentiated together with cell lines from healthy donors across 7 experiments (out of 28). There was no detectable difference in differentiation capacity between healthy and neonatal diabetes lines in these experiments (P>0.05), and cells from both sets of donors overlapped in principal component space (**Fig. S18**). Thus, we included cells from all donors in our analyses irrespective of disease state.

The final merged and QC’ed dataset consisted of 36,044 cells with expression profiles for 11,231 genes (**Fig. S2**).

### Bulk RNA-Seq quality control and processing

Raw RNA-seq data for 546 HipSci cell-lines were obtained from the ENA project: ERP007111 and EGA projects: EGAS00001001137 and EGAS00001000593. CRAM files were merged per cell-line and converted to FASTQ format. Processing of the merged FASTQ files was matched to the single cell processing, as described above. Samples with low quality RNA-seq were discarded based on the following criteria: lines with less than 2 billion bases aligned, with less than 30% coding bases, or with a duplication rate higher than 75%. This resulted in 540 lines for analysis, 108 of which had matched (day0) single cell RNA-seq data available.

Gene-level expression levels were quantified using Salmon, analogously to the alignment, as described for the single cells. Gene expression profiles were normalized using *scran*, to match the single cell data processing, and the *scran* normalized CPM data is log transformed (log2(CPM+1)).

### Variance component analysis

Variance component analysis was performed, per gene, by fitting a random effects model using LIMIX [45] to the gene’s expression profiles across cells. To reduce computational cost, we considered a random subset of 5,000 cells. The experiment, day of collection, and cell line identity were each included as random effects. Full variance component results for all genes are provided in **Table S14**.

### Highly variable genes

The top highly variable genes were computed using *scran*’s *trendVar* and *decomposeVar* functions, using a design matrix to correct for the differentiation experiment-specific effects (i.e. treating each experiment as a different batch). At FDR < 1%, this identified 4,546 highly variable genes.

### Pseudotime definition

We used the first principal component calculated based on the top 500 highly variable genes in our set to represent differentiation pseudotime. This component was linearly re-scaled to take values between 0 (the minimum value observed for any cell) and 1 (the highest value observed). For comparison, we considered three alternative methods for defining pseudo time:

i. We considered diffusion pseudotime (DPT) [46] (**Fig. S7A**). The underlying diffusion map was generated using 15 nearest neighbours and with gene expression represented by the first 20 PCs across the top 500 most highly variable genes. DPT analysis was carried out using the default settings with Scanpy v1.2.2 [47]. There was a Pearson correlation of 0.82 between DPT and the pseudotime definition we used.
ii. We considered calculating pseudotime by projecting each cell on to the principal curve of the first two principal components of the top 500 most highly variable genes (**Fig. S7B**). Principal curve analysis was performed using the R package *princurve [48]*. There was a Pearson correlation of 0.86 between the principal curve pseudotime and the pseudotime definition we used.
iii. We considered representing pseudotime by the mean expression of the differentiation co-expression module. This gene cluster was enriched for GO terms associated with differentiation including ‘anatomical structure morphogenesis’ (GO:0009653), ‘anterior/posterior pattern specification’ (GO:0009952), and ‘response to BMP’ (GO:0071772) (**Table S9; Fig. S7C**). There was a Pearson correlation of 0.64 between the differentiation co-expression module and the pseudotime definition we used. The lower concordance between pseudotime and this module is consistent with the limited set of genes included - the coexpression module only includes genes upregulated during differentiation, and therefore uses no information from changes in expression of pluripotency-associated genes.

### Definition of mesendoderm and definitive endoderm populations

The stage labels post iPSC (mesendo and defendo) were defined using a combination of differentiation stages obtained using the single-cell defined pseudotime and knowledge based on canonical marker genes. Cells were assigned to the mesendo stage if they were collected at day1 or day2, and had pseudotime values between 0.15 and 0.5, corresponding to a pseudotime window around the peak expression of Brachyury (*T*), a marker of mesendoderm (**Fig. S8A**). Cells were assigned to the defendo stage if they were collected at day2 or day3, and had pseudotime values higher than 0.7, corresponding to a pseudotime window with maximal expression of *GATA6*, a marker of definitive endoderm (**Fig. S8B**). Cells with intermediate pseudotime (between 0.5 and 0.7) mostly came from day2, and were not assigned to any stage for the purposes of the initial stage QTL mapping (results shown in Fig. 2). Overall, we assign 28,971 (80%) cells to any of the stages (iPSC, mesendo, defendo).

### Identification of genetic and molecular markers for differentiation efficiency

Differentiation efficiency for each cell line was defined as its average pseudotime across cells at day3, quantified for each experiment and unique donor. To test for associations with molecular markers, we considered stage-specific gene expression levels, again quantified for each donor and experiment (as log2(CPM + 1)).

Three sets of tests were performed. In each case, models were fitted using the lme4 package in R [49], and significance was determined by the Likelihood ratio test. The tested model was:

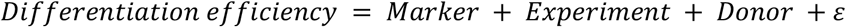

Where Experiment is a random effect grouping sets of samples from the same experiment, and Donor is a random effect grouping samples from the same donor (and cell line). Two sets of Markers were tested - genetic markers (i.e. eQTL SNPs), and expression markers (i.e. expression levels in the iPSC stage/day0), and are presented in **Table S6**, **Table S7**, respectively. For genetic markers, tests were limited to the lead eQTL variant per eGene and differentiation stage.

Genetic markers were validated using data from independent differentiations of individual cell lines. Here, the percentage of CXCR4+ on day 3 (as measured by FACS) was used as a measure of differentiation efficiency, with the following model:

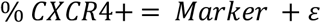

Two sets of tests were performed: (1) all 5 associations (FDR 20%) were tested using data from the original set of cell lines; (2) the strongest association, with the eQTL variant for *DPH3*, was tested using data from new cell lines selected according to their genotype at this locus.

Expression markers were validated by comparison to bulk RNA-sequencing at the iPSC stage (day0). In particular, we tested the association between gene expression in the same cell lines, assayed in separate experiments by bulk RNA-seq of iPSCs, with differentiation efficiency in our experiments, using the model:

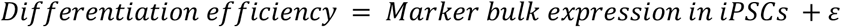

Results of the replication p-values and directions of effect are provided in **Table S7**.

To evaluate whether donor sex had a significant effect on differentiation, we fit the following linear mixed model:

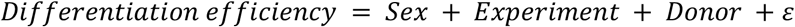

In this model Sex was modelled as a fixed effect and tested for significance using likelihood ratio test, and Experiment and Donor were modelled as random effects, as above.

### *cis* eQTL mapping

A consistent eQTL mapping strategy was applied to bulk RNA-seq expression and expression traits derived from scRNA-seq. We considered common variants (minor allele frequency > 5%) within a *cis-*region spanning 250kb up- and downstream of the gene body for *cis* QTL analysis. Association tests were performed using a linear mixed model (LMM), accounting for population structure and sample repeat structure (see below) as random effects (using a kinship matrix estimated using PLINK [50]). All models were fitted using LIMIX [45]. The values of all features were standardized and the significance was tested using a likelihood ratio test (LRT). To adjust for global differences in expression across samples, we included the first 10 principal components calculated on the expression values in the model, as covariates. In order to adjust for multiple testing, we used an approximate permutation scheme, analogous to the approach proposed in [51]. Briefly, for each gene, we generated 1,000 permutations of the genotypes while keeping covariates, kinship, and expression values fixed. We then adjusted for multiple testing using this empirical null distribution. To control for multiple testing across genes, we then applied the Storey procedure [52]. Genes with significant eQTL were reported at an FDR< 10%.

### Mapping cis eQTL across three stages of differentiation from scRNA-seq data

To map eQTL based on scRNA-seq profiles, we quantified average gene expression profiles (log2(CPM + 1)) across cells for each (donor, day of collection, experiment) combination. This approach retains differences across experiments and days, for cells from the same donor, and is enabled by the pooled experimental design. Accounting for population structure using a kinship matrix is especially important in this context, since aggregated expression values for the same donor from different experiments are essentially replicates and hence genetically identical. We separately mapped eQTL for each differentiation stage (i.e. iPSC, mesendo, defendo), yielding 1,833 (10,840 tested), 1,702 (10,924 tested) and 1,342 (10,901 tested) genes with an eQTL respectively (FDR<10%). eQTL results are provided in **Table S3**).

For comparison, we performed analogous QTL analyses using all cells from day1, and day3 instead of the pseudo-time based differentiation stages. This approach resulted in 1,181 (10,787 tested) and 631 (10,765 tested) genes with an eQTL at day 1 and 3 respectively (**Table S5**).

### Mapping dynamic eQTL (visualisation purposes only)

We performed eQTL mapping across a sliding window on pseudotime, considering bins that contain 25% of all cells, sliding along the pseudotime by a step of 2.5% of cells (Fig. 4A, top middle panel). Similarly to the approach taken for eQTL analysis in individual differentiation stages, expression values are averaged by (donor, day, experiment) combinations, within each window.

### Mapping *cis* eQTL in iPSCs with bulk RNA-seq

To perform *cis*-eQTL mapping in the bulk RNA-seq data, we considered cell lines that had been used to map iPSC eQTL from the scRNA-seq data (bulk data was available for 108 donors out of the 112 day0 single cell donors), and tested the same set of genes. This yielded 2,908 significant genes at an FDR of 10% (out of 10,736 genes tested).

To compare the iPSC eQTL maps derived from bulk and single-cell RNA-seq data, we assessed the nominal significance (P < 0.05) as well as the consistent direction of effect of single-cell iPSC eQTL lead variants (top variant per gene) in the full set of results from the bulk iPSC eQTL analysis and vice versa.

### SNP tagging

We used LD tagging to account for linkage disequilibrium (LD) effects that might cause false positive lead switches and to identify links between GWAS implicated variants and eQTL. To this end, we calculated the LD between lead eQTL variants and either GWAS variants or other eQTL lead variants, using both the 1000 genomes phase3 reference panel and the HipSci dataset to calculate LD between SNPs, taking the union of both sets.

### Lead switching event quantification

Lead switching events were defined as two or three distinct variants that were identified at distinct differentiation stages, found to be significantly associated (FDR < 10%) with the same genes, and that were not in LD (r^2^ < 0.2).

### GWAS Tagging

We performed GWAS tagging using an LD threshold of r^2^ > 0.8. We considered all GWAS variants from the GWAS catalog as available as part of ENSEMBL 94 [53], for all traits and diseases. This analysis was restricted to variants that reached genome-wide significance (P < 5e-8) for any of the traits.

### Allele-specific expression quantification

Duplicated reads were removed from the STAR alignments using Picard Tools (http://broadinstitute.github.io/picard). ASE was quantified at the gene level relative to a heterozygous eQTL lead variant. As a result, for a given eQTL, ASE was only quantified across cells from donors heterozygous for that eQTL variant. This was done following five steps (see **Fig. S19** for a worked example of one gene in one cell): (1) ASE counts were obtained using GATK tools v3.7 in ASEReadCounter mode, with the settings “-minDepth 1 --minMappingQuality 10 −minBaseQuality 2 -rf DuplicateRead”. ASE of a SNP in a given cell was quantified if (i) the cell was heterozygous for that SNP, based on the known donor genotypes, and (ii) the SNP was located in an exonic region (ENSEMBL 75 annotation, as above). The output from GATK tools gives the number of reads mapping to the alternative and reference alleles for each heterozygous SNP in each cell. (2) For each cell, ASE quantifications for each SNP were converted from “alternative allele reads” to “chrB allele reads” using the known phase (indicated as chrA|chrB, where 0=reference, 1=alternative) of each SNP in each donor (e.g. for a SNP with the phase “1|0”, the alternative allele is on chrA, so the number of reads mapping to chrB = number of reference allele reads = total number of reads - number of alternative allele reads). Thus, for each cell, ASE for all SNPs was quantified relative to the genotypes of the chromosomes of that individual, rather than to “reference” or “alternative” alleles. (3) Aggregation of ASE from SNP-level to gene-level. For each gene, this was done by summing the “chrB allele reads” and “total reads” across all SNPs contained in the exons of that gene (as described in the ENSEMBL 75 GTF file). (4) Conversion of quantifications from “chrB allele reads” to “reads from the chromosome containing the alternative allele of the eQTL SNP”, again by using the available phasing information. For each eQTL (i.e. each gene-SNP pair), this provides a consistent definition of ASE across all cells heterozygous for the eQTL SNP (i.e. across cells from multiple donors). Donors that are not heterozygous at the eQTL variant of interest were not used for quantification. (5) Conversion to allelic fractions i.e. quantifications express the allelic reads as a fraction of the total number of reads.

### ASE association tests with cellular factors

ASE quantifies the relative expression of one allele over the other. If one of these alleles is more responsive to a particular environmental factor (e.g. because of preferential transcription factor binding), then ASE is expected to vary systematically with that factor. This observation has previously been used to identify GxE interactions in gene expression across individuals [23]. Here, we applied similar concepts to single-cell RNA-seq, testing for the influence of cellular environmental factors (i.e. cellular processes) on ASE in individual cells. Importantly, these ASE tests are “internally matched”, as potentially confounding batch effects and technical variation affect both alleles in each cell similarly.

Five sets of tests were performed, in a linear modelling framework (Fig. 5, **S13; Tables S13**):

1. Linear pseudotime (“*pseudo*”) tests. The ASE of each gene-eQTL pair was tested for association with pseudotime, across all cells in which ASE was quantified for that pair:

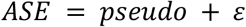
2. Quadratic pseudotime tests. As (1), but with linear pseudotime as a covariate:

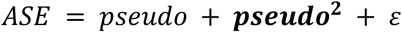
3. Linear cellular factor test. As (1), but with each of 4 cellular factors (“*factor*”) (respiratory metabolism, sterol biosynthesis, G1/S transition and G2/M transition):

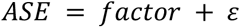
4. Pseudotime-corrected linear cellular factor test. As (3), but with pseudotime included as a covariate:

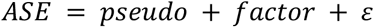
5. Combined pseudotime-factor test. As (4), but testing for the additional effect of (pseudotime x factor) included as a covariate:

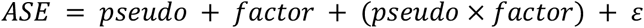

In each case, tests were only performed for eQTL for which ASE was quantified in at least 500 cells. Tests were performed using the statsmodels package in Python (likelihood ratio test). Multiple testing correction was performed independently for each of the five sets of tests, using Benjamini-Hochberg correction.

### Binning ASE across pseudotime

For visualizing ASE as a function of pseudotime or other cellular factors, we averaged ASE across bins of 25% of cells, as done for the sliding window eQTL analysis (above). For each (eQTL x bin) combination, the mean ASE, number of cells, standard deviation, and standard error of the mean (SEM) was calculated (noting that, while each bin contains an equal number of cells, not all cells have quantified ASE for each gene). For each eQTL, to calculate the dynamics of allelic expression across pseudotime (i.e. the expression of transcripts from the ALT and REF chromosomes, as plotted in Fig. 4C), two calculations were performed. First, the mean expression of each gene across the pseudotime bins was calculated using all cells heterozygous for the eQTL SNP (i.e. the cells in which ASE was quantified). The expression of each allele in each pseudotime bin was then calculated by taking the mean ASE +/- SEM, multiplied by the mean expression of that gene (in CPM) in that bin.

### Coexpression and covariation clustering

Grouping of pseudotime-smoothed gene expression and allele-specific expression (see below) was performed by spectral clustering, as implemented by the Python scikit-learn library (Fig. 4). The negative of the Pearson correlation was used as the dissimilarity metric. A range of cluster numbers were tried, with N = 4 judged to be the most clusters possible before highly correlated pairs of clusters were observed.

Grouping of genes by single-cell co-expression was performed using affinity propagation [54], as implemented by the Python scikit-learn library [55]. The Pearson correlation across all cells was used as the similarity/‘affinity’ metric. The top 8,000 highest expressed genes were included in this clustering (as judged by average expression across all cells). This generated a set of 60 co-expression clusters. GO enrichment of each cluster was performed by Fisher’s exact test in Python using GOATOOLS [56], and results are listed in **Table S9**(FDR 10%).

Exemplar co-expression clusters were selected to represent 4 dimensions of cellular state (Fig 5A): cell cycle G1/S transition (cluster 10), cell cycle G2/M transition (cluster 30), cellular respiration (cluster 0), and sterol biosynthesis (cluster 28). This selection was done according to two criteria: (1) strongest enrichment of relevant GO terms. The co-expression clusters showed the largest overrepresentation of genes for the GO terms ‘G1/S transition of mitotic cell cycle’ (GO:0000082; cluster 10), ‘G2/M transition of mitotic cell cycle’ (GO:0000086; cluster 30), ‘respiratory electron transport chain’ (GO:0022904; cluster 0), and ‘sterol biosynthetic process’ (GO:0016126; cluster 28). (2) *a priori* expectation of sources of cell-to-cell variation. Variation in cell cycle stage is a common feature of single-cell datasets [20], while variation in metabolic state during iPSC differentiation is well known [57].

### ChIP-seq experiments and data processing

ChIP-seq was performed using FUCCI-Human Embryonic Stem Cells (FUCCI-hESCs, H9 from WiCell) in a modified endoderm differentiation protocol to that used for the iPSC differentiations (see details below). Cells were grown in defined culture conditions as described previously [58]. Pluripotent cells were maintained in Chemically Defined Media with BSA (CDM-BSA) supplemented with 10ng/ml recombinant Activin A and 12ng/ml recombinant FGF2 (both from Dr. Marko Hyvonen, Dept. of Biochemistry, University of Cambridge) on 0.1% Gelatin and MEF media coated plates. Cells were passaged every 4-6 days with collagenase IV as clumps of 50-100 cells. The culture media was replaced 48 hours after the split and then every 24 hours.

The generation of FUCCI-hESC lines has been described in [59] and are based on the FUCCI system described in [60]. hESCs were differentiated into endoderm as previously described [61]. Following FACS sorting, Early G1 (EG1) cells were collected and immediately placed into the endoderm differentiation media and time-points were collected every 24h up to 72h. Endoderm specification was performed in CDM with Polyvynilic acid (CDM-PVA) supplemented with 20ng/ml FGF2, 10μM Ly-294002 (Promega), 100ng/ml Activin A, and 10ng/ml BMP4 (R&D).

We performed ChIP as described previously [62]. For ChIP-sequencing, ChIP for various histone marks (H3K4me3, H3K27me3, H3K4me1, H3K27ac, H3K36me3) (see **Table S15** for antibodies) was performed on two biological replicates per condition. At the end of the ChIP protocol, fragments between 100bp and 400bp were used to prepare barcoded sequencing libraries. 10ng of input material for each condition were also used for library preparation and later used as a control during peak calls. The libraries were generated by performing 8 PCR cycles for all samples. Equimolar amounts of each library were pooled and this multiplexed library was diluted to 8pM before sequencing using an Illumina HiSeq 2000 with 75bp paired-end reads.

Reads were mapped to GRCh38 reference assembly using BWA [63]. Only reads with mapping quality score ≥ 10 and aligned to autosomal and sex chromosomes were kept for further processing. Peak calling analysis [64] was performed using PeakRanger [65], and only the peaks that were reproducible at an FDR of ≤0.05 in two biological replicates were selected for further processing. Peak calling was done using appropriate controls with the tool peakranger 1.18 in modes *ranger* (H3K4me3, H3K27ac; ‘-l 316 -b 200 -q 0.05’), *ccat* (H3K27me3; ‘-l 316 -win_size 1000 -win_step 100 -min_count 70 --min_score 7 -q 0.05’) and *bcp* (H3K4me1, H3K36me3; ‘-l 316’). Adjacent peak regions closer than 40 bp were merged using the BEDTools suite [66], and those overlapping ENCODE blacklisted regions were filtered out (ENCODE Excludable Mappability Regions [67]). Finally, peaks were converted to GRCh37 coordinates using UCSC LiftOver [68].

## Data availability

All HipSci data can be accessed from http://www.hipsci.org. Bulk RNA-seq data are available under accession numbers: ERP007111 (ENA project) and EGAS0000100113, EGAS00001000593 (EGA projects). Single cell RNA-seq data for the open access lines (study 3963) are available under the accession numbers ERP016000 (ENA project).

## Supporting information

Supplementary Figures and small Supplementary Tables S5, S11, S15

## Acknowledgements

This work was funded with a strategic award from the Wellcome Trust and Medical Research Council (WT098503). We thank the staff in the Cellular Genetics and Phenotyping and Sequencing core facilities at the Wellcome Trust Sanger Institute. D.D.S. and M.J.B. were supported by fellowships from the EMBL Interdisciplinary Postdoc (EI3POD) program under Marie Skłodowska-Curie Actions COFUND (grant number 664726). The Ludovic Vallier (L.V.) lab is funded by the ERC advance grant New-Chol and the core support grant from the Wellcome Trust and Medical Research Council to the Wellcome–Medical Research Council Cambridge Stem Cell Institute. We thank Yuanhua Huang (Stegle Lab) for assistance in applying the Cardelino software, and Anna Osnato (Vallier Lab) for assistance with ChIP-seq data.

## Author contributions

Wrote the paper with input from all authors - A.C., D.S., J.M., O.S.

Pilot study - M.C., F.B., D.M., A.K., K.N.

Developed the experimental protocol - M.C., J.G.

Experiments - M.C., J.G., I.M., S.A., A.I.

eQTL mapping and analysis - A.C.

scRNA-seq processing and QC - D.M., A.C.

scRNA-seq exploratory data analysis - A.C., D.M., D.S.

Donor mapping - D.M.

Processing of the bulk RNA-seq data and genotype information - M.J.B.

ChIP-seq data analysis - P.M.

Allele-specific expression analysis - D.S.

Differentiation efficiency marker analysis - D.S.

Developed the eQTL mapping approach & pipeline - A.C., M.J.B., D.M., D.S.

Supervised and designed the research - O.S., L.V., J.M., M.C.

