## Supplementary Figures and small Supplementary Tables S5, S11, S15 for "Single-cell RNA-sequencing of differentiating iPS cells reveals dynamic genetic effects on gene expression"

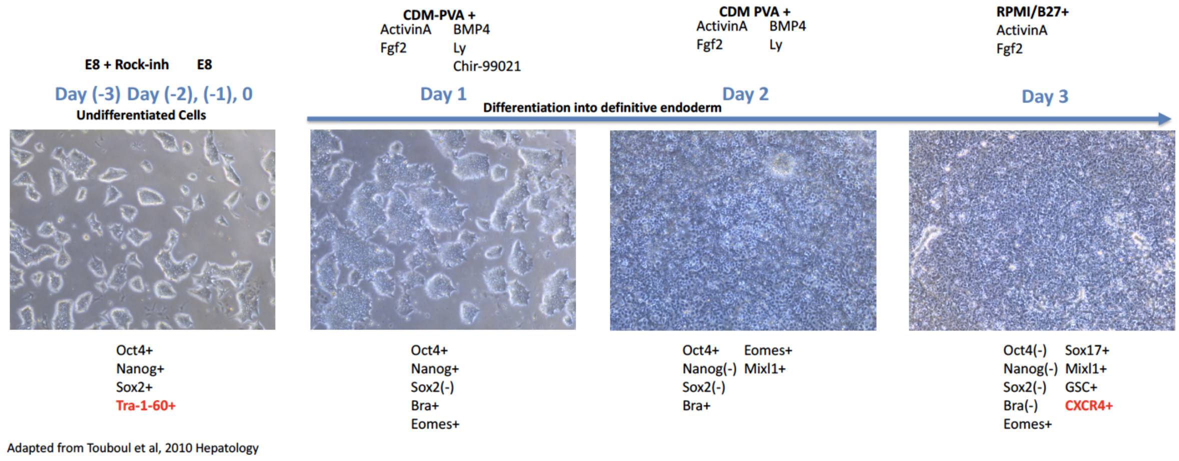


**Figure S1 | Endoderm differentiation protocol.** Schematic representation of the chemically defined protocol used to initiate differentiation towards definitive endoderm (adapted from (Touboul et al. 2010)). Tra-1-60 and CXCR4 are canonical cell surface markers used to sort live cells by differentiation stage.

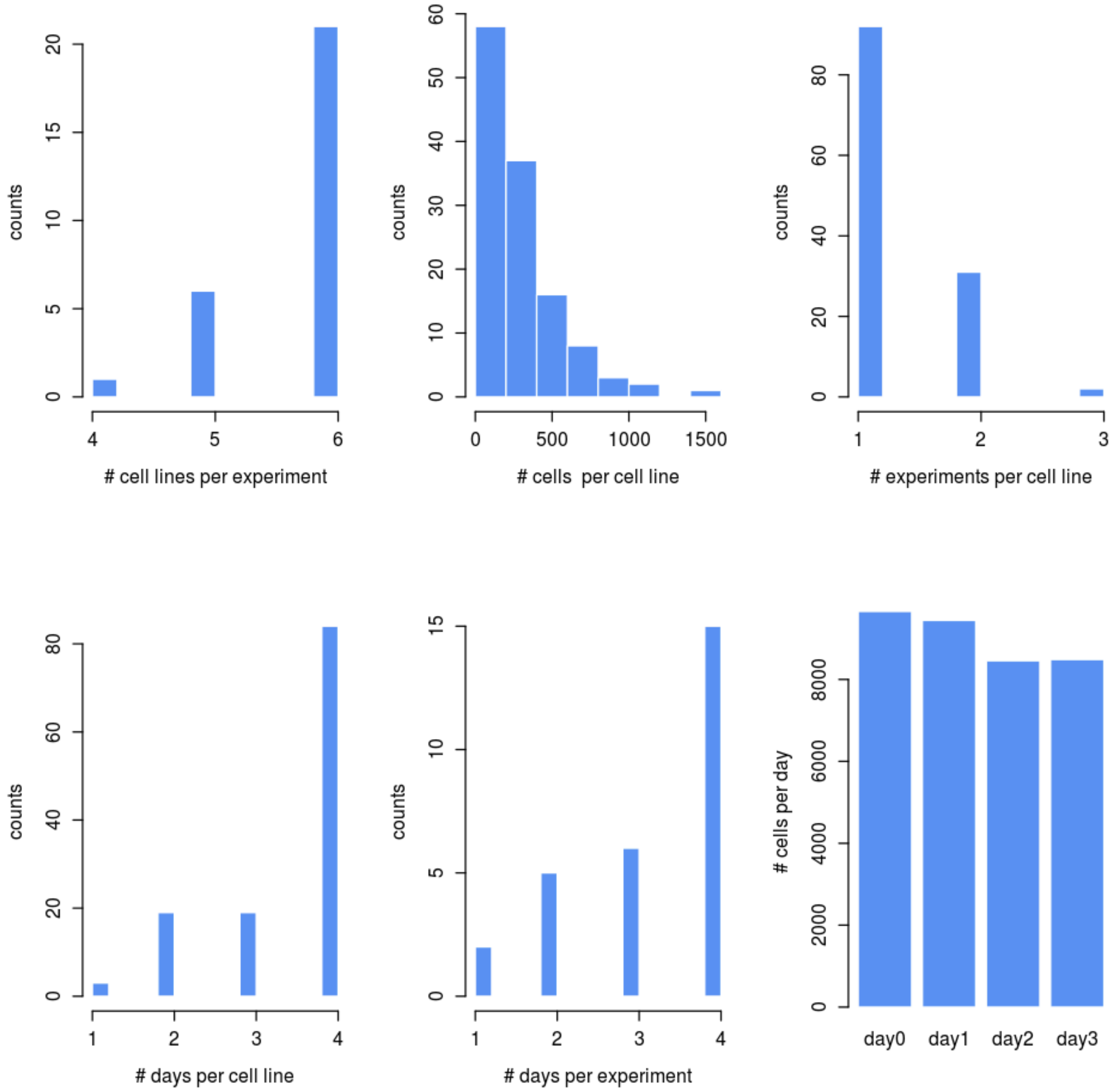


**Figure S2 | Overview of experimental metrics.** Statistics for number of cells, donors, experiments, days, and combinations. Cell counts are shown after quality control.

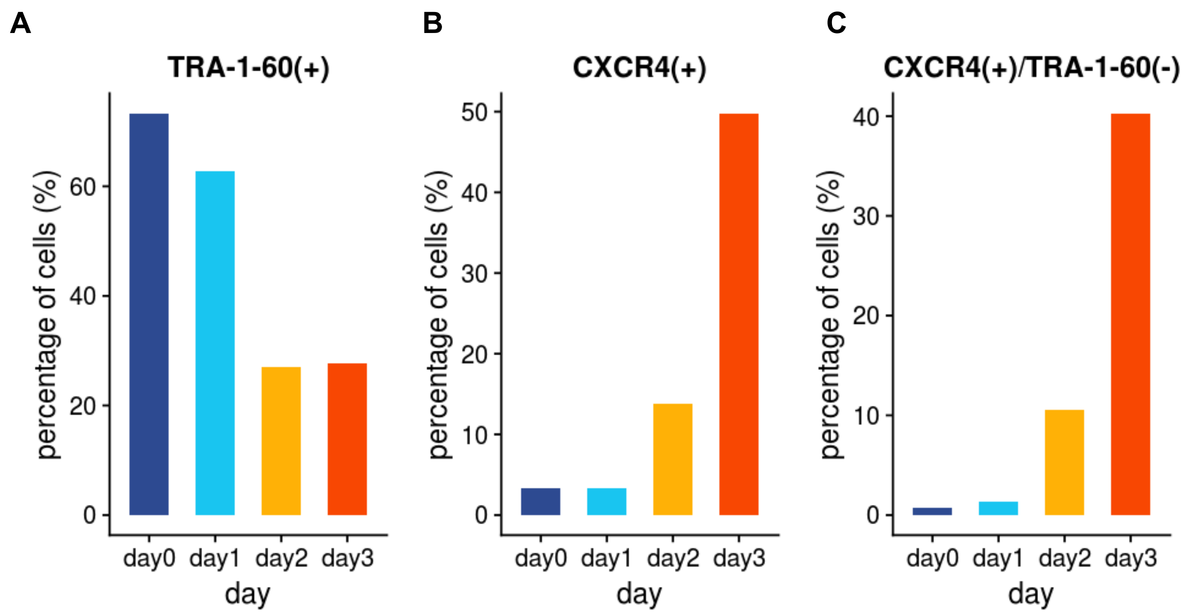


**Figure S3 | Cell surface marker expression across differentiation.** Shown are the percentages of cells that are **(A)** positive for TRA-1-60, a pluripotency marker, **(B)** positive for CXCR4, a definitive endoderm marker, and **(C)** positive for CXCR4 and negative for TRA-1-60, across all cell lines and all experiments.

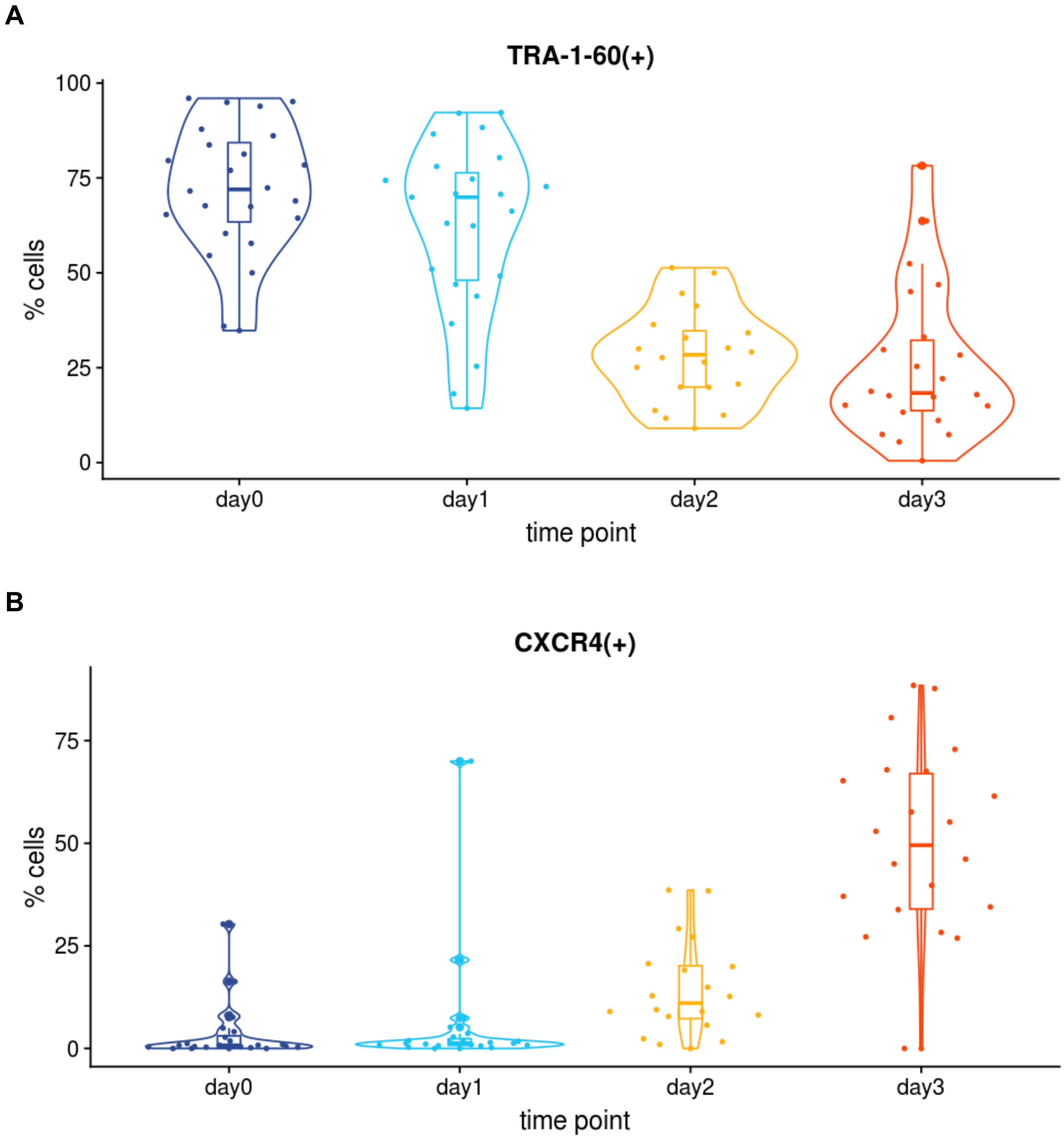


**Figure S4 | Distribution of cell surface marker expression across differentiation experiments.** Shown are the percentages of cells that are (**A**) positive for TRA-1-60, a pluripotency marker, (**B**) positive for CXCR4, a definitive endoderm marker, for each experiment, by day.

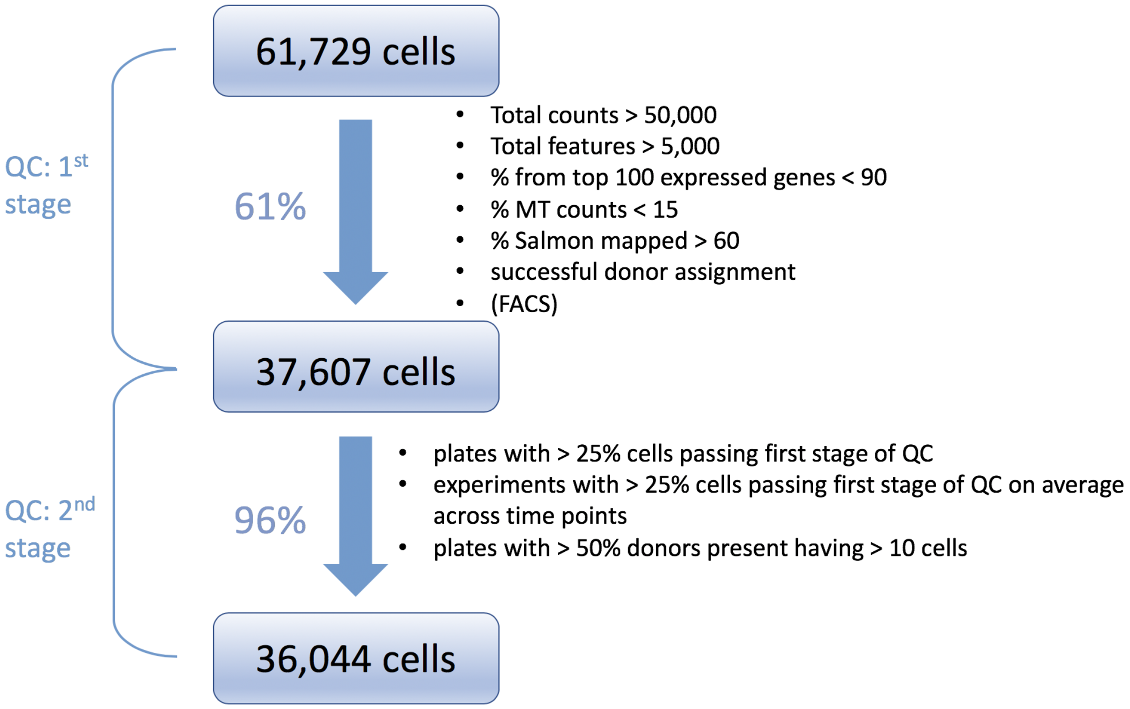


**Figure S5 | Workflow of scRNA-seq quality control.** Quality control (QC) was carried out in two stages. First, QC was applied on the level of individual cells using conventional quality metrics. Second, QC was applied on the level of scRNA-seq processing plates and experimental batches, using aggregate quality metrics to retain cells fom  high-quality plates and experiments. Total numbers of cells before and after each QC step are shown, along with the percentage of cells retained in each QC step.

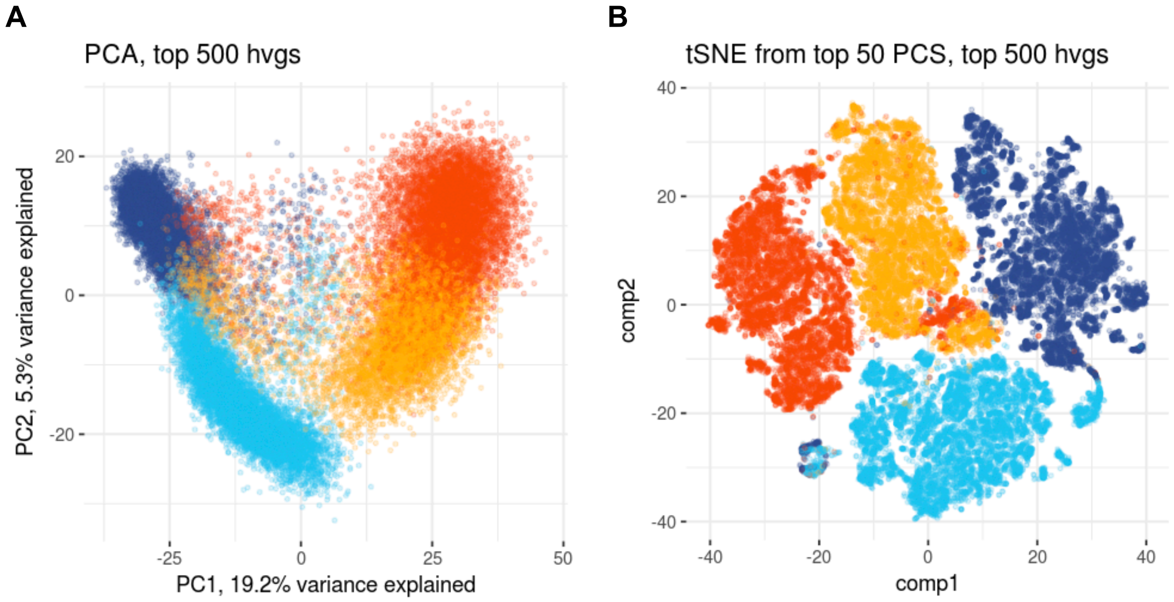


**Figure S6 | Overview of PCA and t-SNE representations of the full scRNA-seq dataset.** (**A**) First two principal components (PC1 and PC2) computed on top 500 highly variable genes (**Methods**). Axes labels show the percentage of variance explained. (**B**) t-SNE plot, computed from the first 50 PCs, on the top 500 highly variable genes.

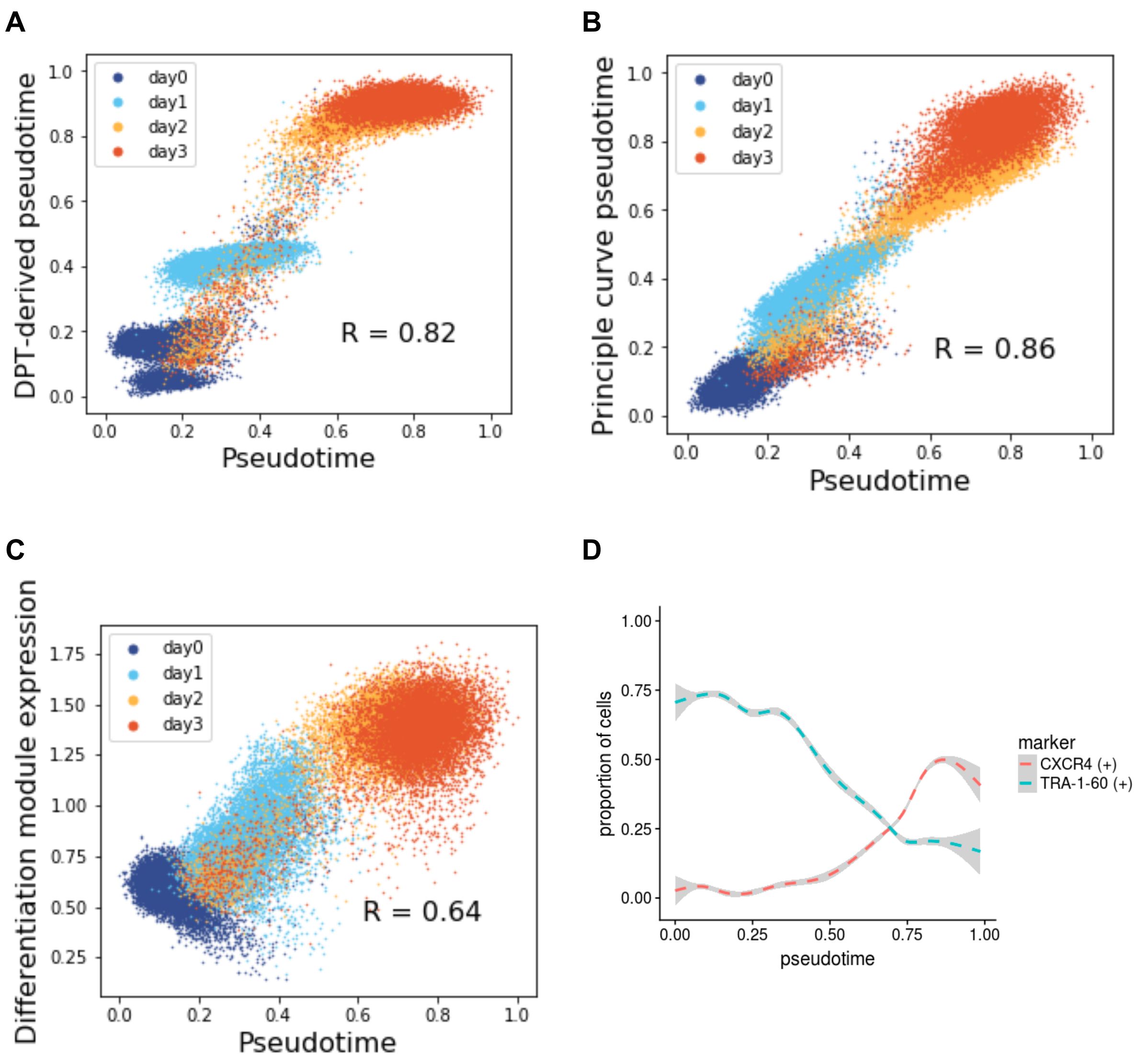


**Figure S7 | Evaluation of pseudotime definition.** (**A**) Comparison of the pseudotime defined based on principal component analysis with diffusion pseudotime (DPT) (Haghverdi et al. 2016). The underlying diffusion map was generated using 15 nearest neighbours and with gene expression represented by the first 20 PCs across the top 500 most highly variable genes (**Methods**). (**B**) Comparison of PCA-based pseudotime with an alternative pseudotime based on projection of each cell on to a principle curve in the first two principal components of the top 500 most highly variable genes (**Methods**). (**C**) Comparison of pseudotime to the mean expression of a set of 124 co-expressed genes that are associated with cell differentiation (**Methods**). (**D**) Scatter plot of FACS markers as a function of the PCA-based pseudotime, showing expected trends.

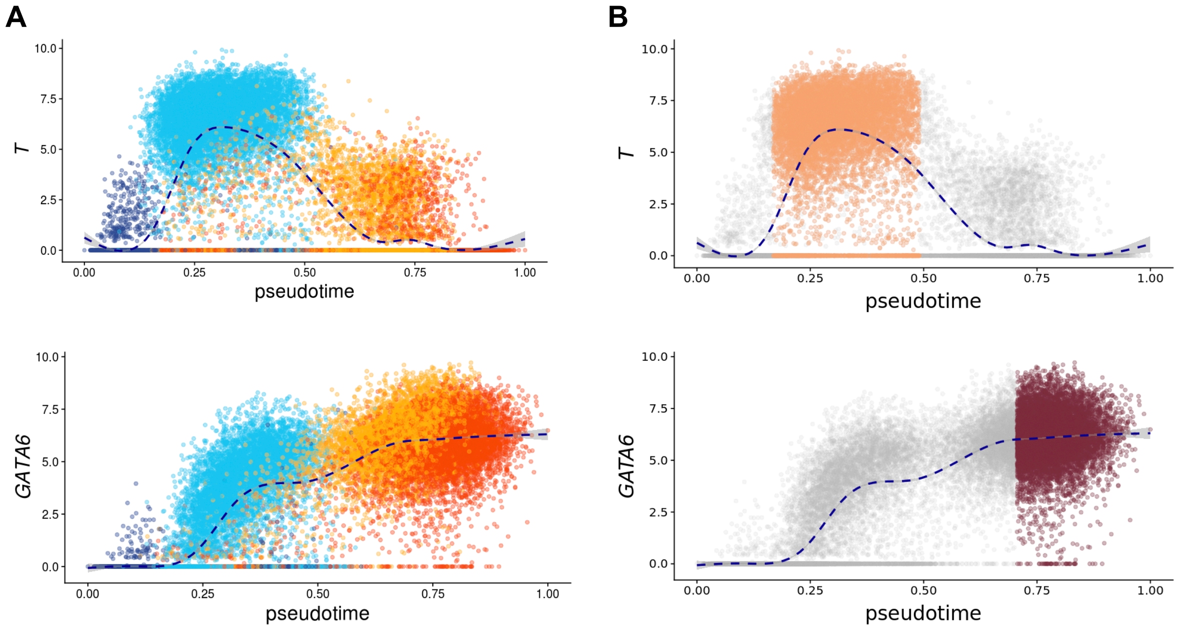


**Figure S8 | Definition of pseudotime-based developmental states.** (**A**) Expression of exemplar canonical markers for mesendoderm (*T*) and definitive endoderm (*GATA6*) along pseudotime. Cells are coloured by the time point of collection, as in **Fig. 1D** (**B**) On the same plots as in **A**, cells assigned to mesendo and defendo, respectively, are highlighted (**Methods**).

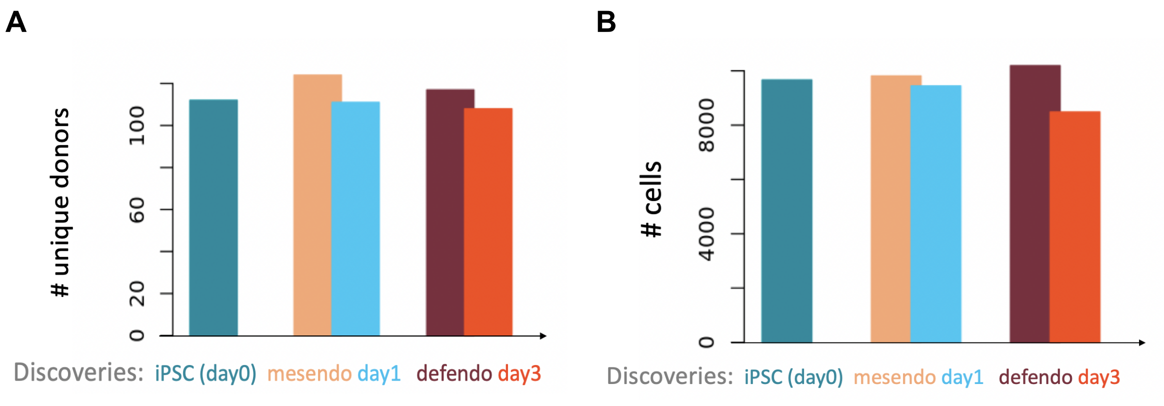


**Figure S9 | Comparison of numbers of donors and cells at each time point and each differentiation stage.** Related to **Fig. 2B**. (**A**) The number of donors for which gene expression data were assayed at day0, day1, and day3, compared to the number of donors in the pseudotime-inferred mesendo and defendo stages. (**B**) As for **A**, with the number of cells.

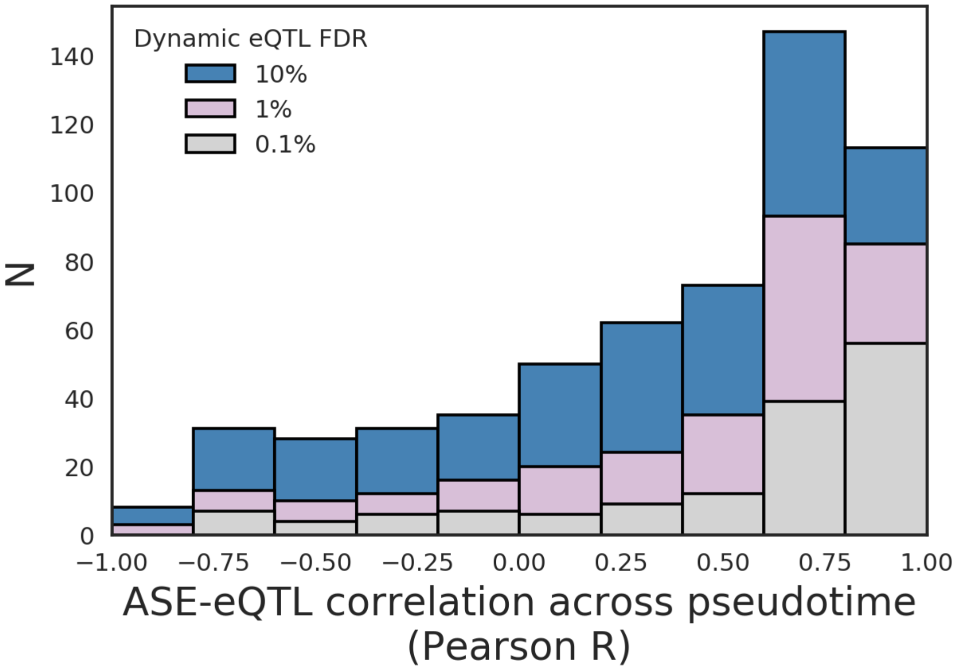


**Figure S10 | Comparison of eQTL effect and ASE dynamics across pseudotime.** The correlation between eQTL effect (i.e. -log10(p) x direction of effect) and ASE across pseudotime, at different FDR thresholds, with 1% FDR corresponding to the set of eQTL plotted in **Fig 4A**.

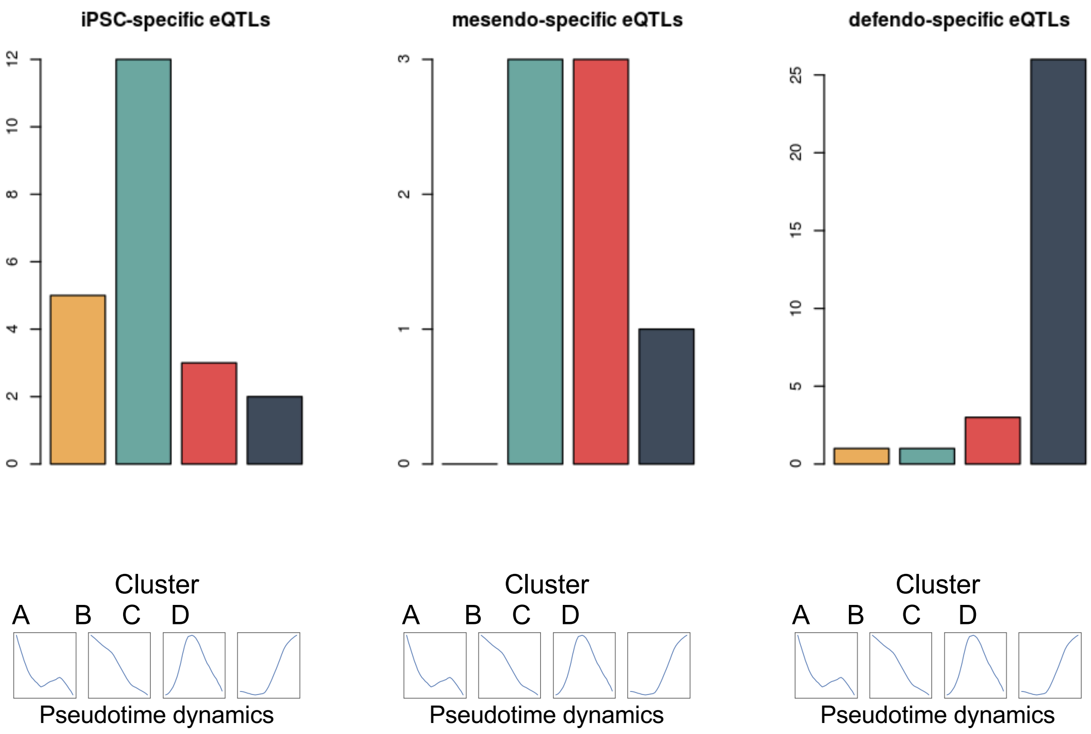


**Figure S11 | Assignment of stage-specific eQTL to dynamic eQTL clusters.** The numbers of each of the 3 classes of stage-specific eQTL (i.e. iPSC-, mesendo-, and defendo-specific eQTL) that are assigned to each of the 4 dynamic eQTL clusters.

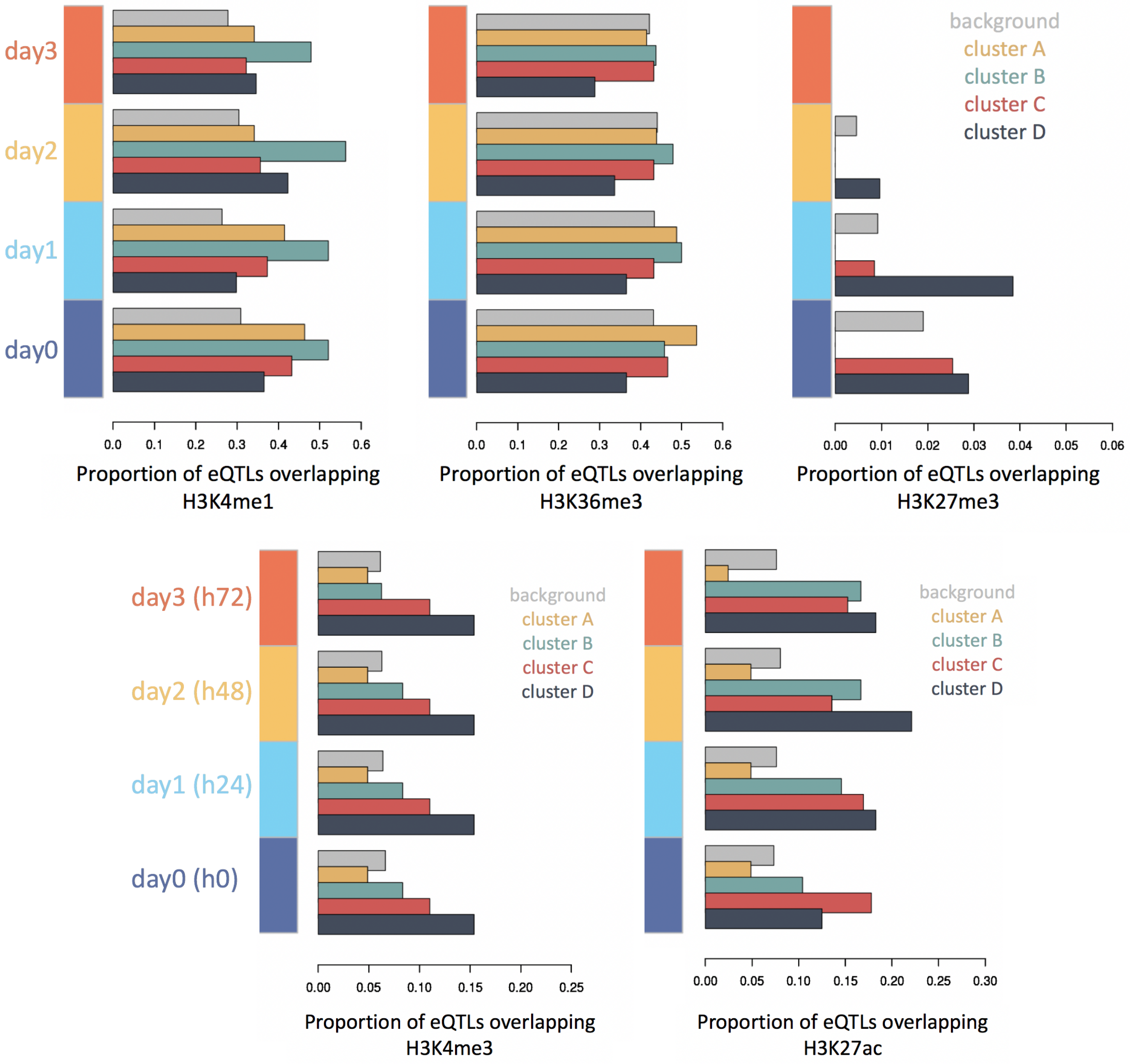


**Figure S12 | Epigenetic marks of dynamically regulated eQTL SNPs across pseudotime dynamics clusters, and time points.** Related to **Fig 4E**. Proportions of dynamic eQTL in each category overlapping each epigenetic mark at each time point are shown. Proportions of overlap with ‘background’ eQTL (i.e. those without an interaction with pseudotime at FDR 1%) are shown in grey for comparison.

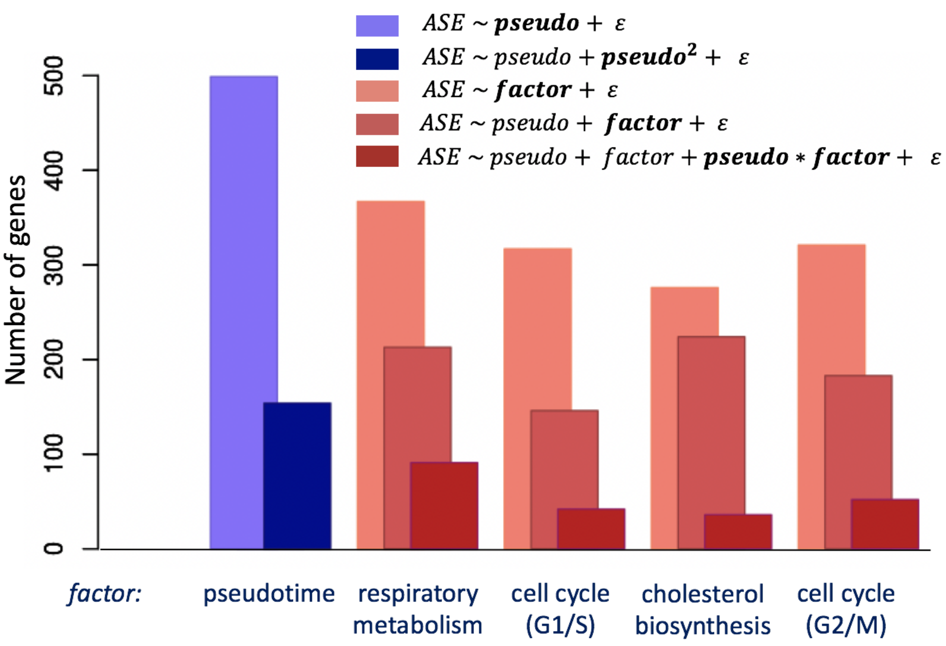


**Figure S13 | Summary of allele-specific expression interaction test results for each tested cellular state.** Results from **Tables S13.** The number of significant interactions in each category are provided. Bars represent the number of genes with at least one eQTL that is significant for each test described in the inset (**Methods**), FDR < 10%.

**A**

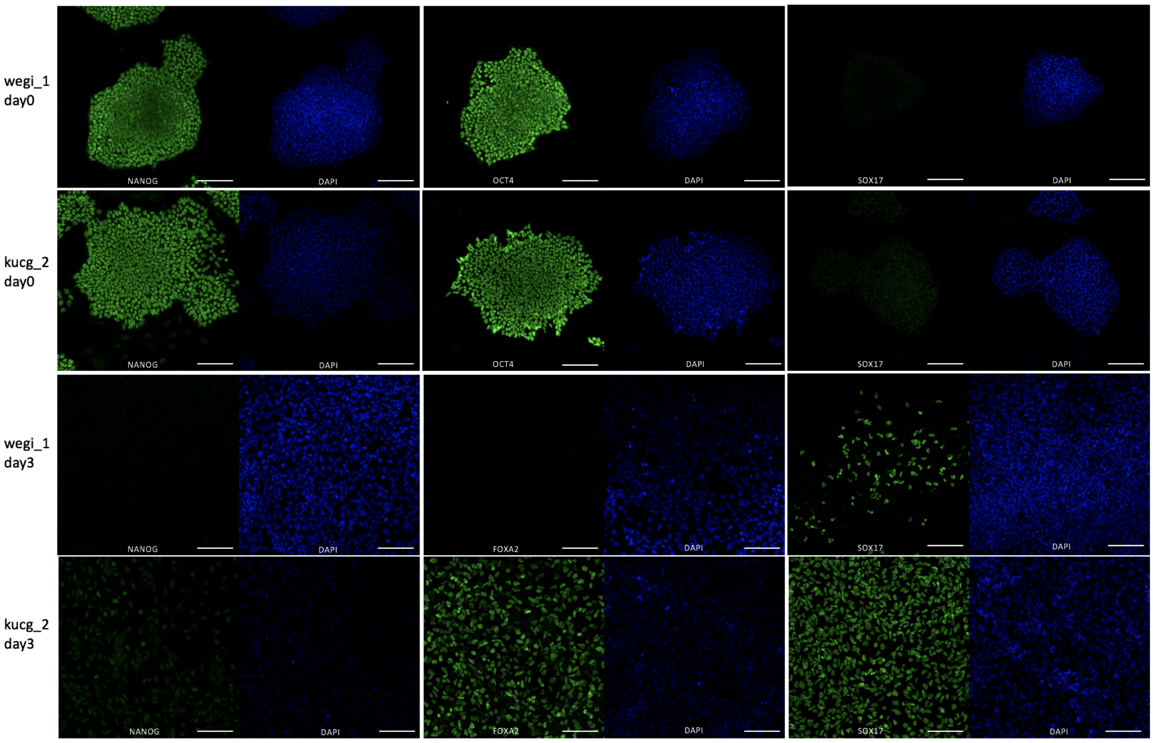

wegi_1
day0
kucg_2
day0
wegi_1
day3
kucg_2
day3
NANOG
DAPI
OCT4
SOX17
FOXA2


**B**

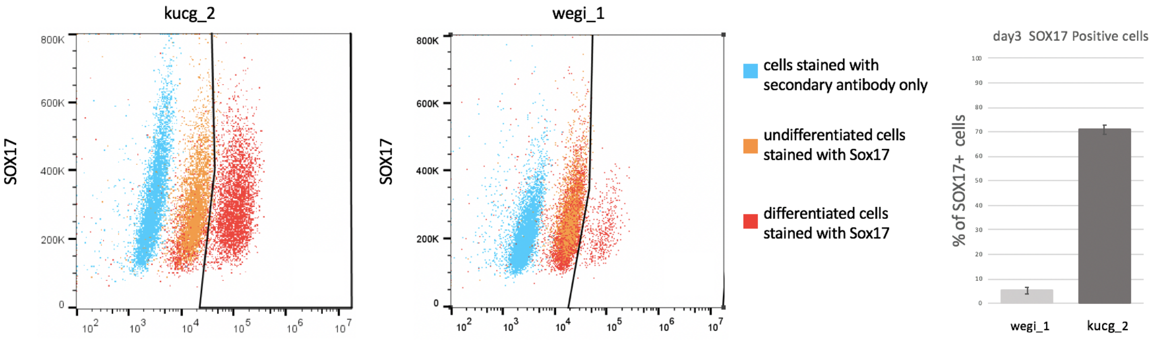

kucg_2
wegi_1
SOX17
800K
600K
400K
200K
0
cells stained with secondary antibody only
undifferentiated cells stained with Sox17
differentiated cells stained with Sox17
day3 SOX17 Positive cells
% of SOX17+ cells
wegi_1
kucg_2

**C**

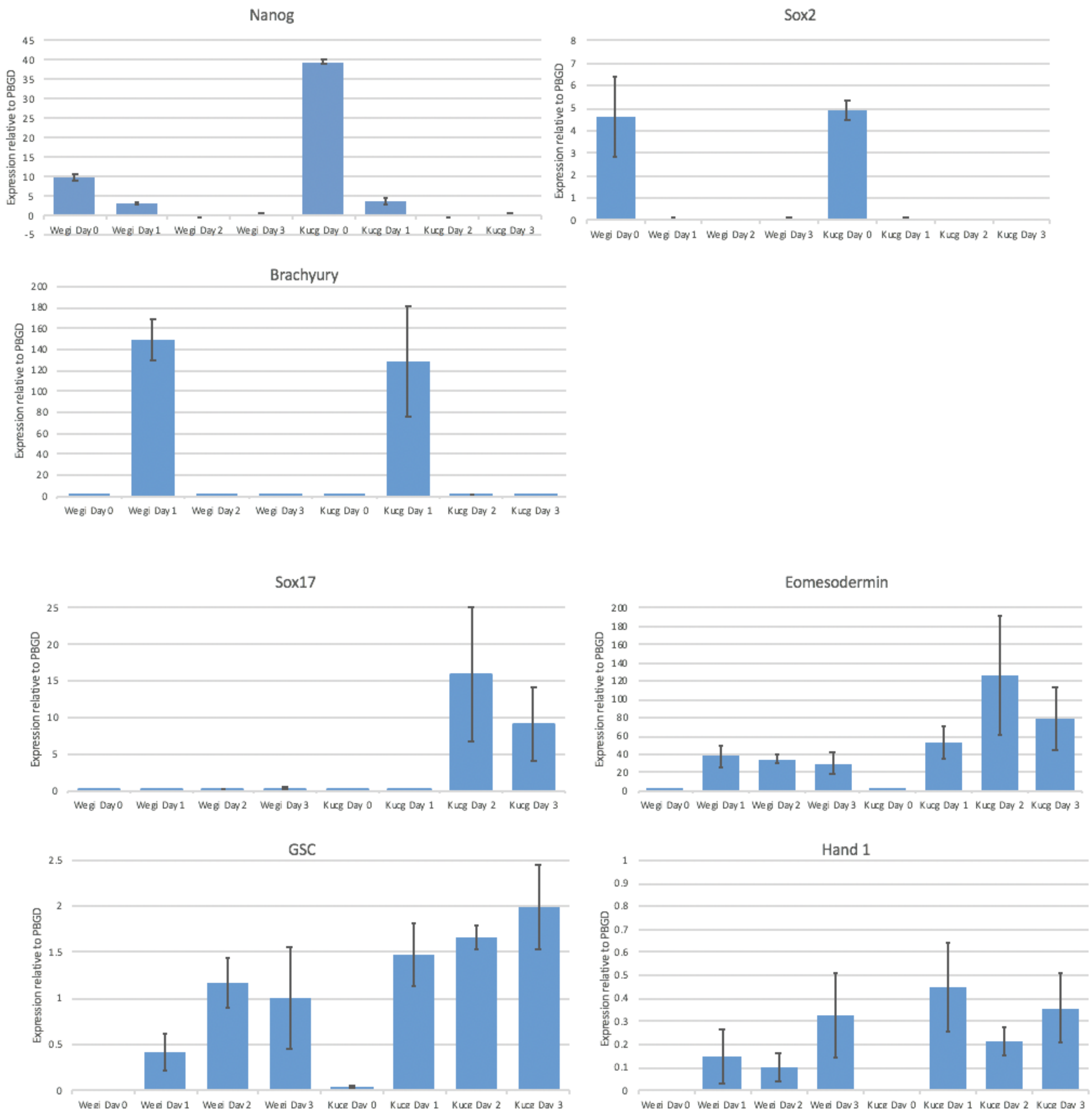


**Figure S14 | Validation of the definitive endoderm differentiation protocol.** Wegi_1 and kucg_2 were identified as poor and highly efficient lines, respectively, for definitive endoderm differentiation. Shown is the expression of various markers in the iPSC and differentiated state as assessed by immunofluorescence (**A**)**,** FACS (**B**), and qPCR (**C**)**.**

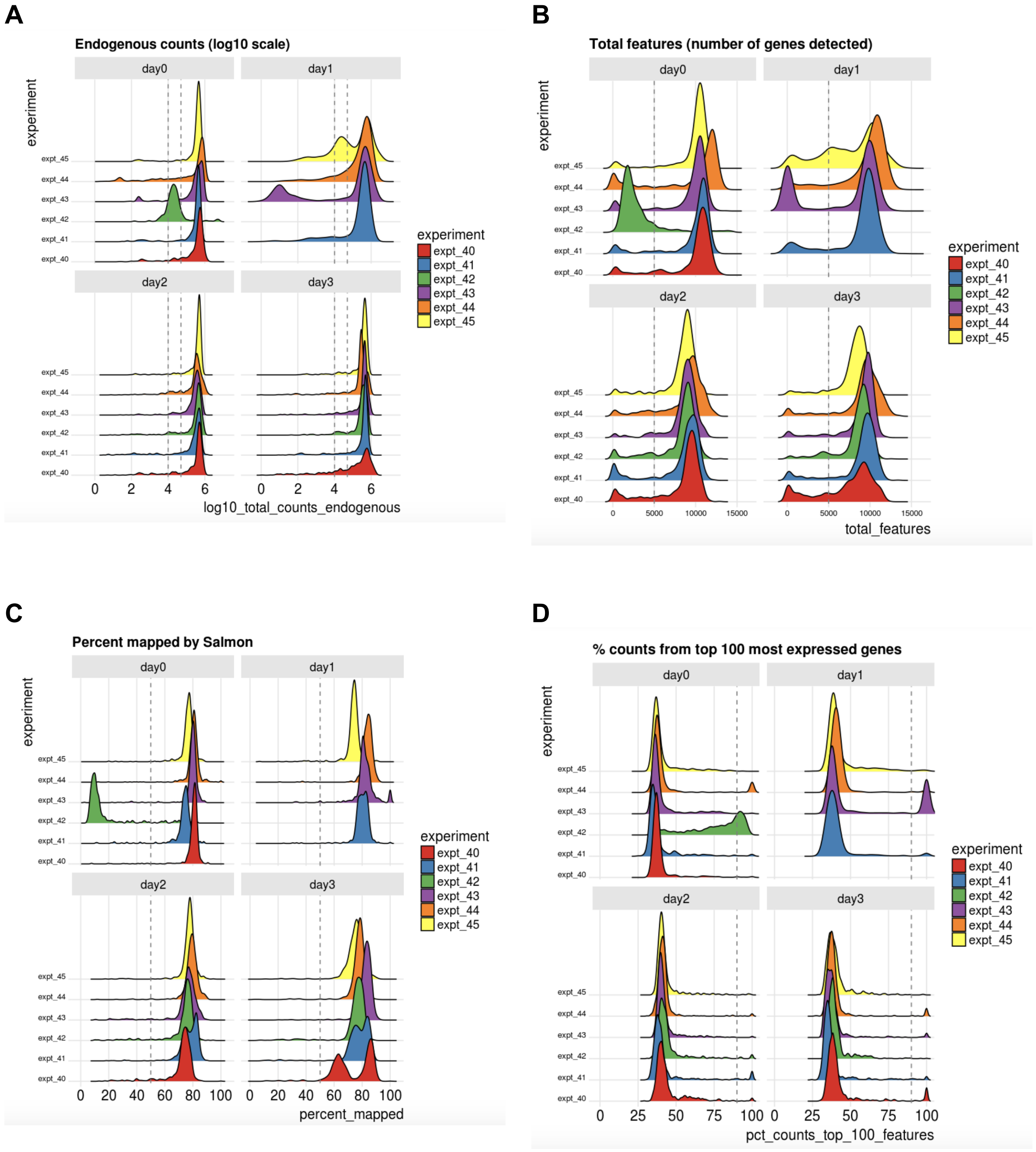
A
Endogenous counts (log10 scale)
day0
day1
day2
day3
experiment
expt_40
expt_41
expt_42
expt_43
expt_44
expt_45
log10_total_counts_endogenous
B
Total features (number of genes detected)
total_features
C
Percent mapped by Salmon
percent_mapped
D
% counts from top 100 most expressed genes
pct_counts_top_100_features

E

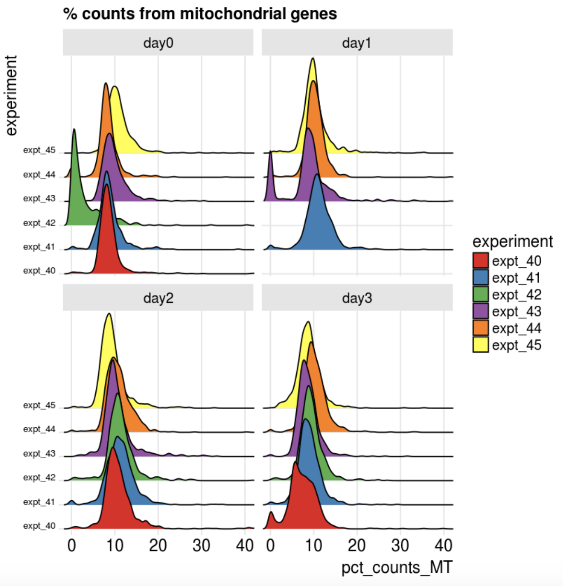


**Figure S15 | Distributions of quality control metrics across all days in an illustrative subset of 6 differentiation experiments**. (**A**) Number of counts for endogenous genes per cell. (**B**) Total number of features (i.e. genes) detected per cell. (**C**) Salmon mapping rate i.e. the percentage of reads successfully mapped to the transcriptome by Salmon. (**D**) Percentages of counts coming from the top 100 most highly expressed genes for each cell. (**E**) Percentage of counts from mitochondrial genes for each cell. In all plots, vertical dashed lines indicate the threshold applied to define the low-quality cells that are excluded from further analysis (**Methods**).

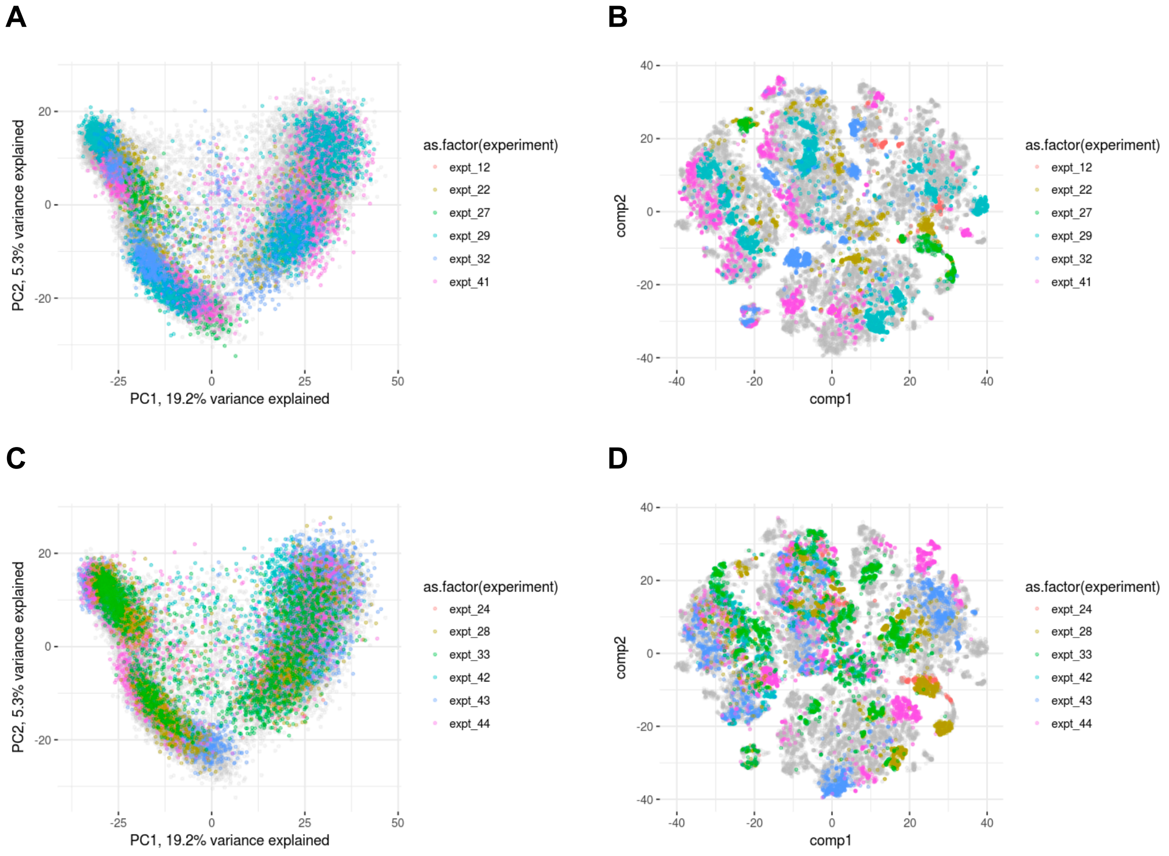


**Figure S16 | Comparisons of gene expression across experiments.** PCA and t-SNE representations of two randomly selected sets of 6 experiments for which data were available across all days. (**A**) PCA plot for the first subset of 6 experiments (colours), against the background of all cells (grey). (**B**) t-SNE plot of the same cells as in **A**. (**C, D**) As for **A, B**, for a different subset of cell differentiation experiments.

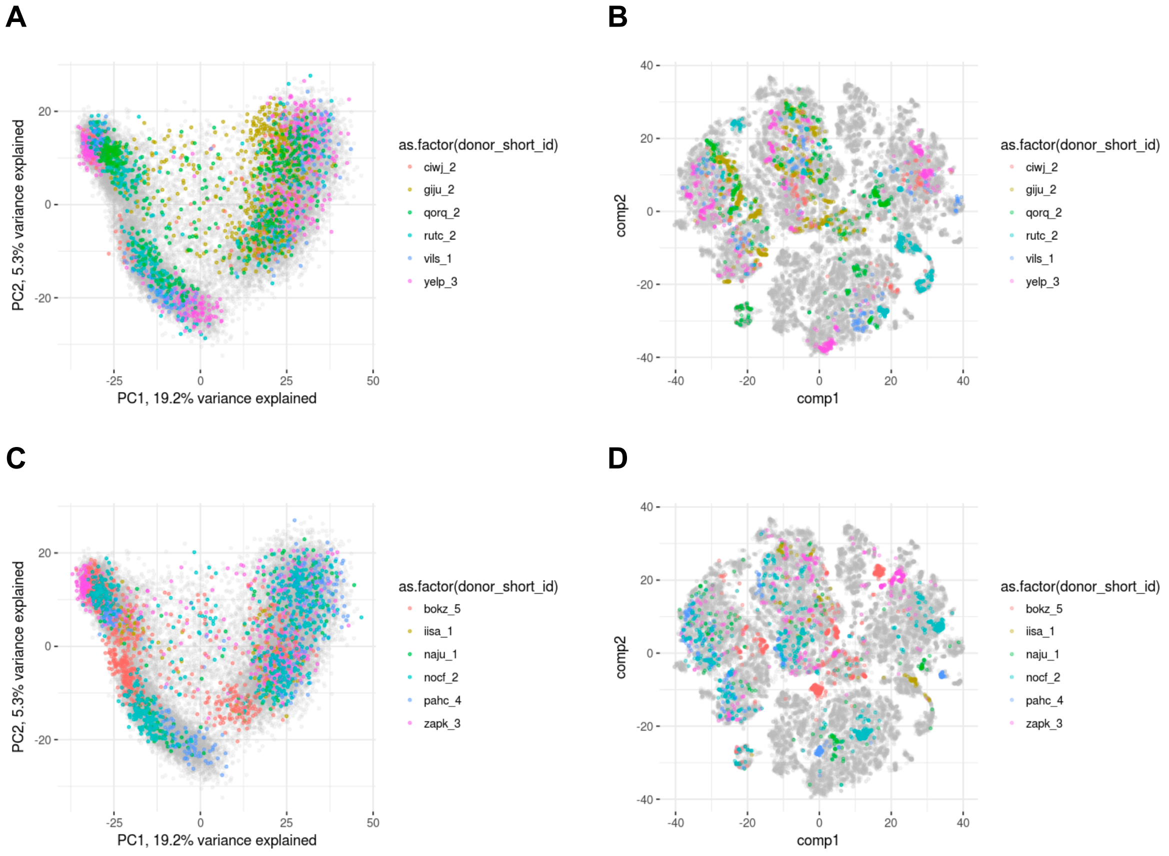


**Figure S17 | Comparison of expression patterns across cell lines.** PCA and t-SNE representations of cells from two randomly selected sets of 6 cell lines. (**A**) PCA plot for the first subset of 6 cell lines (colours), against the background of all cells (grey). (**B**) t-SNE plot of the same cells as in **A**. (**C, D**) As for **A, B**, for a different subset of cell lines.

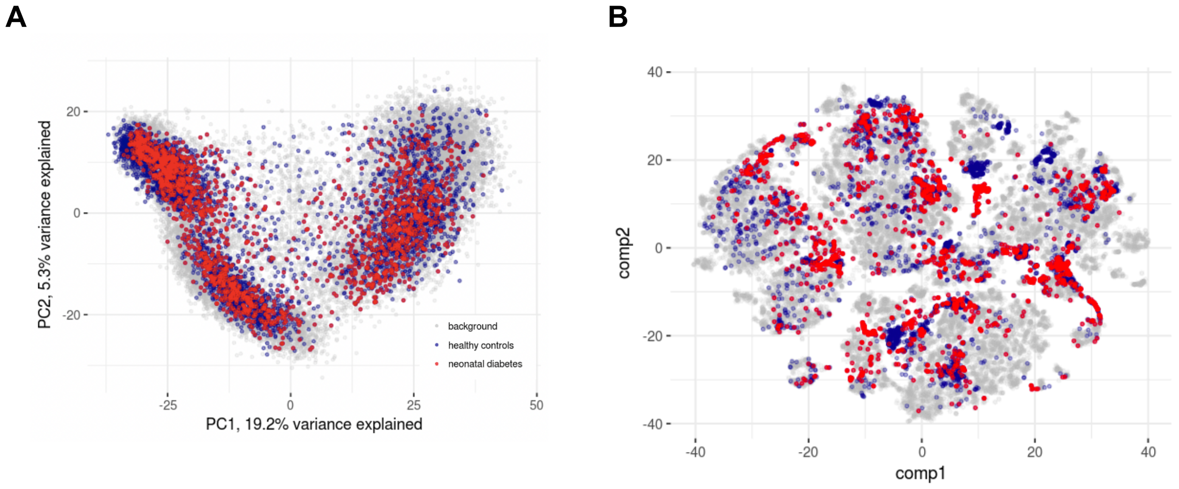


**Figure S18 | Comparison of expression patterns between healthy and diseased cell lines.** PCA and t-SNE representations of cells from neonatal diabetes lines, compared to healthy lines from the same experiments. (**A**) PCA plot for cells from the neonatal diabetes cell lines (red), cells from healthy lines from the same seven experiments (dark blue), against the background of all cells (grey). (**B**) t-SNE plot of the same cells as in **A**.

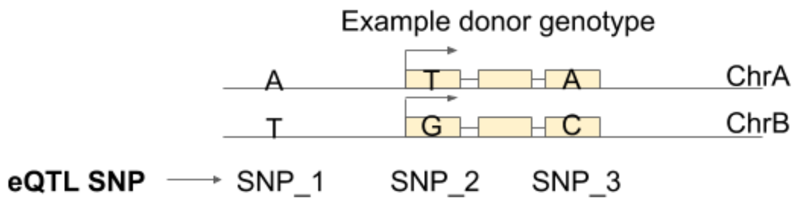


SNP information

| SNP | REF | ALT |
|---|---|---|
| SNP_1 | T | A |
| SNP_2 | G | T |
| SNP_3 | A | C |

Genotype information

| Donor | SNP_1 | SNP_2 | SNP_3 |
|---|---|---|---|
| Example | 1\|0 | 1\|0 | 0\|1 |

1) Count allele-specific reads from RNA-seq

| SNP | REF | ALT |
|---|---|---|
| SNP_2 | 11 | 19 |
| SNP_3 | 24 | 16 |

2) Convert to ChrA/ChrB read counts

| SNP | ChrA | ChrB |
|---|---|---|
| SNP_2 | 19 | 11 |
| SNP_3 | 24 | 16 |

3) Sum counts to gene level

| Gene | ChrA | ChrB |
|---|---|---|
| Example gene | 43 | 27 |

4) Define chromosomes relative to alleles of the eQTL SNP

| Gene | Chr with the REF eQTL allele | Chr with the ALT eQTL allele |
|---|---|---|
| Example gene | 27 | 43 |

5) Convert to allelic fraction

| Gene | Fraction of reads from Chr with the ALT eQTL allele |
|---|---|
| Example gene | 0.61 |

**Figure S19 | Worked example of the ASE quantification procedure.** A toy example is shown, to illustrate the steps involved in quantifying ASE for an eQTL. ASE is first quantified for SNPs, then combined at gene level, then re-defined relative to the genotype and phase of the eQTL variant. **SNP information:** the REF and ALT alleles. **Genotype information:** the genotype of each individual, including phasing information, in “chrA|chrB” format, where 0 is REF and 1 is ALT (e.g. “0|1” indicates chrA is the REF allele and chrB is the ALT allele).

### Supplementary Tables

**Supplementary Table S1. Differentiation experiment metadata.** This table is supplied as an external data file.

**Supplementary Table S2. Cell line and donor metadata.** This table is supplied as an external data file.

**Supplementary Table S3. Summary of single-cell eQTL results**, at all stages. All lead eQTL SNP-gene pairs are provided. This table is supplied as an external data file with fields defined below.

| Table field | Description |
|---|---|
| ensembl_gene_id | Ensembl ID (Ensembl version 75) |
| snp_id | Lead variant, SNP ID in the format [chromosome]_[position]_[reference]_[alternative allele] |
| p_value | Nominal P-value |
| empirical_feature_p_value | Gene-level corrected P-value using 1,000 permutations |
| global_corr_p_value | Q-value, globally corrected P-value using Storey procedure |
| beta | Effect size of the eQTL |
| beta_se | Standard error of the effect size |
| gene_name | HGNC symbol |
| snp_chromosome | Variant chromosome |
| snp_position | Variant position |
| ref_allele | Variant reference allele |
| alt_allele | Variant alternative allele |
| stage | Stage at which the eQTL was discovered |
| stage_specific | Whether the eQTL is specific to the stage in which it was discovered (True/False) |
| interaction_qtl | Whether the eQTL is found to be sensitive to any of the measured cell states, including pseudotime (True/False) |
| dynamic_qtl | Whether the eQTL is found to be sensitive to pseudotime (True/False) |
| in_HipSci | Whether the eQTL is tagging an iPSC eQTL from (Mirauta et al., 2018) |

| | |
|---|---|
| n_gtex_tissues | How many GTEx tissues is the eQTL tagging (0-49) |
| GWAS_tagging | Whether the eQTL is tagging a GWAS variant |

**Supplementary Table S4. Summary of bulk iPS eQTL results.** The list of genes with a significant (FDR < 10%) eQTL. This table is supplied as an external data file with fields defined below.

| Table field | Description |
|---|---|
| ensembl_gene_id | Ensembl ID (Ensembl version 75) |
| snp_id | Lead variant, SNP ID in the format [chromosome]_[position]_[reference]_[alternative allele] |
| p_value | Nominal P-value |
| empirical_feature_p_value | Gene-level corrected P-value using 1,000 permutations |
| global_corr_p_value | Q-value, globally corrected P-value using Storey procedure |
| beta | Effect size of the eQTL |
| beta_se | Standard error of the effect size |
| gene_name | HGNC symbol |
| snp_chromosome | Variant chromosome |
| snp_position | Variant position |
| ref_allele | Variant reference allele |
| alt_allele | Variant alternative allele |

**Supplementary Table S5. Summary of the type and number of eQTL.** Including all eQTL discovered based on single cell (at iPS, mesendo, defendo stage, and day0, day1, day3 time point) and bulk (only iPS) RNA traits. Shown are the number of genes that were considered for QTL mapping, as well as the number of genes for which a QTL was detected.

| | Number of genes with an eQTL (FDR < 0.1) | Number of genes tested | Number of cells in pool | Sample size (number of donors) | Number of (donor, day, experiment) combinations |
|---|---|---|---|---|---|
| bulk iPS | 2,908 | 10,736 | - | 108 | - |
| sc iPS (day0) | 1,833 | 10,840 | 9,661 | 111 | 136 |
| sc mesendo | 1,702 | 10,924 | 9,809 | 123 | 224 |
| sc defendo | 1,342 | 10,901 | 10,187 | 116 | 238 |
| sc day1 | 1,181 | 10,787 | 9,443 | 111 | 138 |
| sc day3 | 631 | 10,765 | 8,485 | 108 | 127 |

**Supplementary Table S6. Associations between eQTL variants and differentiation progress.** Related to **Fig. 3B**. Results of association tests between identified eQTL (iPSC, mesendo,defendo) with differentiation progress. Coefficients and p-values of the tests are provided. This table is supplied as an external data file.

**Supplementary Table S7. Associations between discovered marker genes and differentiation progress.** Related to **Fig. 3C**. Results of association tests between the 38 significantly associated genes (FDR < 10%) ("candidate_marker_gene") and differentiation progress. Coefficients and nominal p-values for all tests are provided. The column heading suffix ("_all_lines", "_female_lines", "_male_lines") indicates the set of cell lines in which the association test was performed. The chromosome on which each gene is located is also provided. This table is supplied as an external data file.

**Supplementary Table S8. Coexpression clusters.** List of genes (HGNC symbols) and their corresponding coexpression cluster. This table is supplied as external data file.

**Supplementary Table S9. Gene ontology (GO) enrichments for all clusters** (Fisher's exact test). Column key: 'cluster_label': cluster label, 'GO': GO term ID number, 'NS': GO term category, 'enrichment': whether it is an enrichment (e) or depletion (p), 'name': full name of the GO term, 'ratio_in_study': ratio of proteins in the cluster that are annotated with this GO term, 'ratio_in_pop': ratio of all proteins that are annotated with this GO term, 'p_uncorrected': raw p-value from Fisher's exact test, 'depth': depth of GO term in the GO tree, 'study_count': number of proteins in the cluster annotated with this GO term, 'p_fdr_bh': p-value after correction for multiple testing by Benjamini-Hochberg. This table is supplied as an external data file.

**Supplementary Table S10. Enrichments of transcription factor binding in coexpression clusters.** The ChEA 2016 database (Lachmann et al. 2010) was used to identify transcription factor target genes.

**Supplementary Table S11. Functional annotation of clusters.** See **Tables S9,S10** for supporting GO and ChIP-seq enrichment data.

| Cluster label | Functional annotation |
|---|---|
| 0 | Respiration |
| 10 | G1/S transition |
| 28 | Sterol biosynthesis |
| 30 | G2/M transition |

**Supplementary Table S12. GWAS tagging results.** For the joint set of eQTL identified at iPSC, mesendo, defendo ($r^2$> 0.8). This table is supplied as external data file.

**Supplementary Table S13. GxE results by ASE analysis.**
**S13a**. Results from linear test as described in (1,3) from **Methods,** *ASE association tests with cellular factors.*
**S13b.** For pseudotime only, including quadratic pseudotime (2).
**S13c.** For all factors, including pseudotime as a covariate (4).
**S13d.** Non linear interactions, including pseudotime and another factor (5). These tables are supplied as external data files with fields defined below.

| **Table field** | **Description** |
|---|---|
| ensembl_gene_id | Ensembl ID (Ensembl version 75) |
| snp_id | Lead variant, SNP ID in the format [chromosome]_[position]_[reference]_[alternative allele] |
| pval | Nominal P-value |
| coef | Interaction effect size |
| ncells | Number of cells considered for ASE |
| index | Unique eQTL (SNP-gene pair) identifier in the format ([ensembl_gene_id], [snp_id]) |
| mean_ase | Average ASE |
| factor (**a,b,c**) or factor1 & factor2 (**d**) | Factor tested (those can be pseudotime, G2_M_transition, sterol_biosynthesis, respiration, G1_S_transition) |

**Supplementary Table S14. Variance component results.** Related to **Fig. 1B**. The variance components of cell line, experiment, and time point are provided.

**Supplementary Table S15. Antibodies used for ChIP-seq experiments.**

| **Antibody raised against** | **Catalogue number** | **Company** |
|---|---|---|
| Histone H3 | ab1791 | Abcam |
| Histone H3 (tri methyl K4) | ab8580 | Abcam |
| Histone H3 (tri methyl K27) | C15200181<br>(MAb-181-050) | Diagenode |
| Histone H3 (mono methyl K4) | ab8895 | Abcam |
| Histone H3 (acetyl K27) | ab4729 | Abcam |
| Histone H3 (tri methyl K36) | ab9050 | Abcam |
